# Single cell analysis of RA synovial B cells reveals a dynamic spectrum of ectopic lymphoid B cell activation and hypermutation characterized by NR4A nuclear receptor expression

**DOI:** 10.1101/2021.05.14.443150

**Authors:** Nida Meednu, Javier Rangel-Moreno, Fan Zhang, Katherine Escalera-Rivera, Elisa Corsiero, Edoardo Prediletto, Edward DiCarlo, Susan Goodman, Laura T Donlin, Soumya Raychauduri, Michele Bombardieri, Costantino Pitzalis, Dana E Orange, Accelerating Medicines Partnership Rheumatoid Arthritis and Systemic Lupus Erythematosus (AMP RA/SLE) Network, Andrew McDavid, Jennifer H Anolik

## Abstract

Ectopic lymphoid structures (ELS) can develop in rheumatoid arthritis (RA) synovial tissue, but the precise pathways of B cell activation and selection are not well understood. Here, we identified a unique B cell population in the synovium characterized by co-expression of a family of orphan nuclear receptors, NR4A1 (also known as NUR77), NR4A2 (NURR1) and NR4A3 (NOR1), that is highly enriched at both early and late stages of RA. NR4A B cells are rare in healthy peripheral blood, RA blood, and SLE kidney, but share markers with blood transcriptomic signatures that peak during RA disease flare. Using combined single cell transcriptomics and B cell receptor (BCR) sequencing, we demonstrate that NR4A synovial B cells have an activated transcriptomic profile that significantly overlaps with germinal center (GC) light zone (LZ) B cells and an accrual of somatic hypermutation that correlates with loss of naïve B cell status. NR4A B cells uniquely co-express lymphotoxin *β* and IL6, supporting important functions in ELS promotion and pro-inflammatory cytokine production. Further, the presence of shared clones in this activated B cell state and NR4A expressing synovial plasma cells (PC) and the rapid up-regulation with BCR stimulation points to *in situ* differentiation. Taken together, we identified a dynamic progression of B cell activation in RA synovial ELS, with NR4A transcription factors having an important role in antigen activation and local adaptive immune responses.

**One sentence summary:** B cells in the rheumatoid arthritis synovium undergo a spectrum of in situ activation, with the NR4A family of transcription factors having an important role in antigen stimulation, local adaptive immunity, and pathological B cell responses.

## INTRODUCTION

Rheumatoid arthritis is a chronic autoimmune disease characterized by inflammation of the synovial tissue leading to joint damage and disability. Although dramatic advances in treatment options for RA have been achieved over the last two decades, a significant number of RA patients do not achieve remission or low disease activity (1, 2), highlighting the need for new therapies and biomarkers of response. B cells play a key role in RA disease pathogenesis both through autoantibody mediated and antibody independent functions and are key treatment targets in the disease (3–6). Though B cell clonal expansion is detected in RA blood (7) and synovium (8), the mechanisms and location of antigen specific B cell priming remains unclear.

B cell aggregates are present in the RA synovium (9, 10) at both early and late stages of disease and have been linked to important clinical outcomes including treatment response, bone erosion, and radiographic progression (3, 10). Ectopic lymphoid structures (ELS) are found in at least 40% of RA patient synovia with 10-25% of those structures displaying features of a functional germinal center (GC) (11–14). ELS organization can recapitulate the architecture of secondary lymphoid organs (SLO), with areas that resemble dark zones (DZ) (proliferating and AID expressing B cells) and areas that resemble light zones (LZ) (rich in follicular dendritic cells and T cells in addition to B cells) (15). ELS in RA synovial tissue has been shown to support antigen-driven selection and differentiation of autoreactive B cells, facilitating diversification, somatic hypermutation (SHM) (14) and class switching (16) for in situ production of autoantibodies (17), suggesting that these structures are functional. Many critical factors driving ectopic lymphoid neogenesis, including lymphoid chemokines, are shared with factors driving SLO development, but the cellular components that produce these key factors may differ and are the subject of ongoing investigation. In mouse colitis lymphotoxin (LT) expressing B cells can support ELS formation and lead to severe inflammatory disease (18). The role of B cells during ELS in RA synovial tissue, however, is unclear and which B cell subsets may be crucial for this process is still unknown.

Single cell analyses of RA synovial tissue recently identified four B cell states including naïve, memory B cells, age-associated B cells, and PCs (19). To improve our understanding of the cellular, transcriptional, and antibody repertoire dynamics during human B cell activation in ELS, here we performed unbiased single-cell transcriptomic and repertoire profiling of RA synovial B cells. These single-cell antibody repertoires paired with single-cell transcriptomics allowed us to define novel transcriptional B cell states, resolving functional B cell heterogeneity and gene expression dynamics. We identified an abundant population of synovial B cells characterized by up-regulation of NR4A1-3 along with other immediate early response genes (EGR1/3, FOS, JUN) and activation markers (CD69, CD83). These cells show evidence of SHM, class-switching, and clonal relationship with a subset of NR4A expressing PCs. Gene expression profiling of synovial NR4A B cells demonstrated enrichment for genes associated with GC LZ B cells and expression of elevated levels of chemokine receptors and chemotactic factors involved in ELS, including LT-*β* and IL6. This subset of B cells is enriched in synovial tissue and found at very low levels in peripheral blood and another autoimmune target tissue, the kidney in lupus nephritis. Notably, RA synovial B cells spontaneously express NR4A1 protein by flow cytometric and histologic analysis, and NR4A cluster genes are enriched in synovial samples with a lymphoid histologic pathotype (9). In an RA flare cohort (20), an NR4A B cell transcriptomic signature peaked in the blood during flare. Finally, stimulation of normal B cells through the BCR *in vitro* up-regulated NR4A1-3. Together our data support *in situ* activation of B cells with accrual of SHM in synovium ELS and NR4A as a read-out of antigen stimulation, local adaptive immunity, and pathological B cell responses.

## RESULTS

### Unbiased scRNA-seq analysis of RA B cells reveals multiple distinct subsets including a unique NR4A B cell population in the synovium

In order to define the diversity of B cells in the synovium, we isolated B cells from lymphoid-rich RA synovial tissues and paired blood and performed single cell transcriptomic and BCR sequencing on the 10X Chromium platform (Fig. 1 and table S1). The mean frequency of B cells by flow cytometry was 12.65% (2.7%-21.6%, n=4) of CD45^+^ synovial cells. The majority of synovial B cells were IgD^-^, whereas 80% of blood B cells were naïve (IgD^+^CD27^-^). We also observed synovial enrichment of CD24^-^CD27^hi^ plasmablasts/plasma cells (1.9%-7.6%) (fig. S1B).

**Fig 1.**
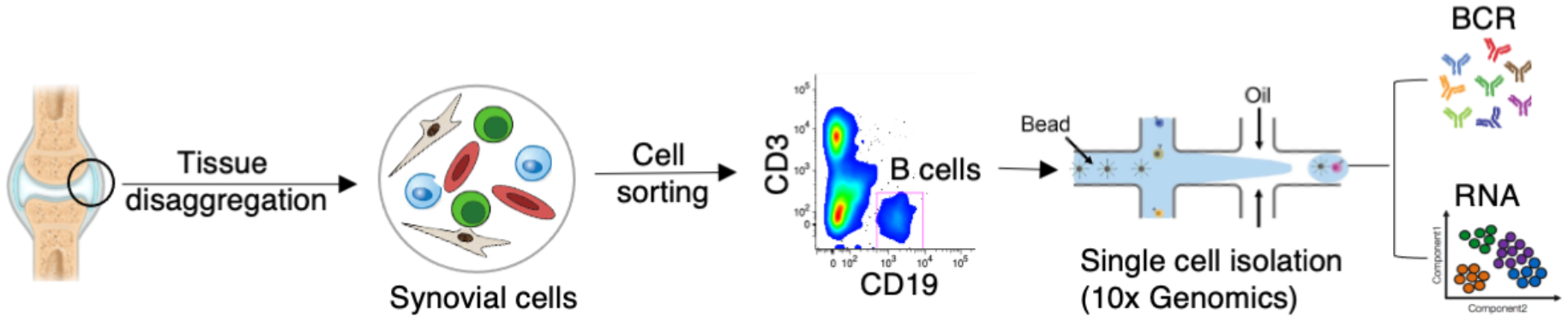
Overview of work flow. Synovial tissues were disaggregated using a validated protocol (66) and B cells were sorted by flow cytometry cell sorting. We then performed scRNA-seq on sorted tissue and blood B cells using the 10x genomic platform with poly-A selected, 5’ initiated expression and V(D)J libraries generated from each single cell.

We sequenced 4,115 B cells (2,589 synovial and 1,526 peripheral blood) with an average median genes per cell of 1,427 (755-1,868). After exclusion of cells with high mitochondrial RNA content, dimensionality reduction using t-distributed stochastic neighbor embedding (t-SNE) (21) and unsupervised clustering using Seurat package (22) on the combined data set of 3,786 cells, 8 conserved B cell subsets or clusters were defined. Differentially expressed genes (DEGs) were identified by calculating the difference between the average expression by cells in the subset versus cells not in the subset (Fig. 2). Two subpopulations of plasma cells (PC(i) and PC (ii)) were characterized by high expression of transcription factors required for PC differentiation and maintenance (*XPB1, IRF4* and *PRDM1)* and immunoglobulin genes (Fig. 2, A-B and D). Expression of *IGHD* and *TCL1A* identified three naïve B cell subpopulations (Naïve(i), Naïve (ii) and Naïve (iii)) (Fig. 2, A and B). Two B cell subpopulations expressed high levels of *CD27, HOPX*, *S100A10 and S100A4*, markers associated with memory B cells (23–26). The LMNA^+^ designated subpopulation also uniquely expressed the *LMNA* gene, not previously reported in memory B cells (Fig. 2, A-B and D). The final subpopulation was labeled NR4A^+^ as it expressed high levels of the nuclear family receptor 4 (*NR4A1, NR4A2, NR4A3*). This subpopulation also showed high expression of markers associated with activation and GC formation like *CD69*, *CD83* and *GPR183*, the latter a molecule mediating B cell migration in lymphoid follicles (27, 28) (Fig. 2, A-B and D).

**Fig 2.**
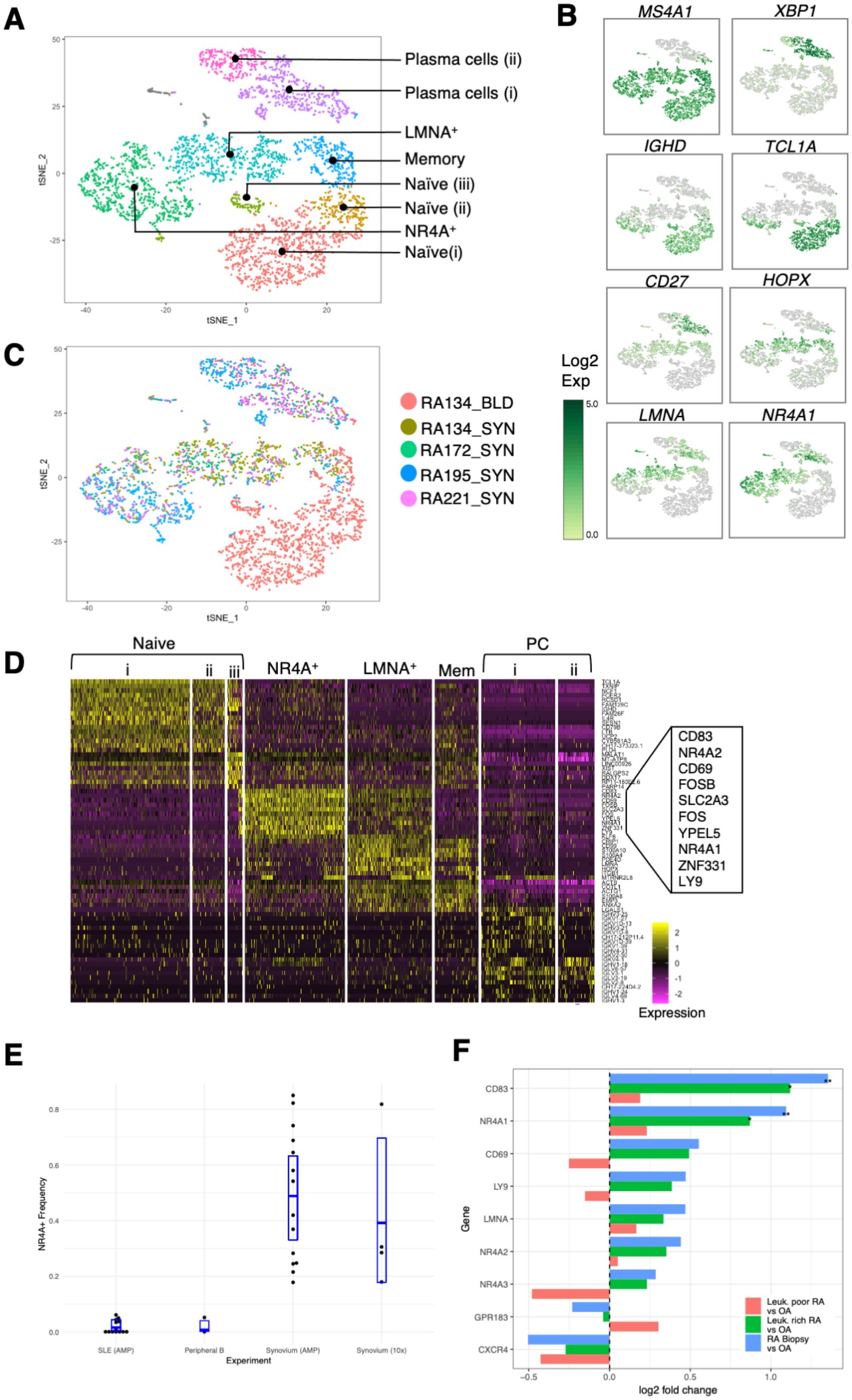
Single-cell RNA sequencing identifies an enrichment of B cells expressing nuclear orphan family receptors (NR4As) in RA synovial tissue. (**A**) tSNE visualization of 8 B cell clusters from 3,786 B cells from 4 RA synovial tissues (SYN) and one paired blood sample (BLD). (**B**) Markers identified 2 clusters of plasma cells, 3 clusters of naïve, a memory, LMNA^+^ and NR4A^+^ cluster. (**C**) The same tSNE map as in (A) with cells labeled by sample ID. (**D**) Heatmap displays top differentially expressed genes (DEGs) in each cell clusters. Top 10 DEGs of NR4A^+^ cluster are magnified. (**E**) Box plots display frequency of NR4A^+^ B cells calculated using SingleR. Single-cell RNA sequencing of Synovium (AMP) (n=14) from Zhang et al, 2019 (19) (ImmPort SDY998), SLE (AMP)(n=13) from Arazi et al, 2019 (30) (ImmPort SDY997), peripheral B cells from Zheng et al, 2017 (31) and RA134_BLD and current study (Synovium (10x)). Horizontal blue line is the mean and the blue box represent 90% CI. (**F**) Bar graphs plotting log2 fold change of NR4A^+^ cluster genes in leukocyte poor RA, leukocyte rich RA or RA biopsy vs. OA using bulk RNA sequencing from Zhang et al, 2019 (19) (ImmPort SDY1299). * significant at 5% and ** significant at 1%

The NR4A^+^ and LMNA^+^ subpopulations were restricted to synovial samples. Naïve B cells were the main population in the blood, whereas the memory and PC subpopulations were present in both blood and synovial tissue (Fig. 2C, and fig. S2C).

### Unique enrichment of NR4A B cells in RA synovium compared to blood and other tissues

Using a supervised classification method SingleR (29), we next examined the presence of NR4A B cells in an independent RA synovial tissue cohort and other tissues where single cell data was available. In SLE kidney (30) and peripheral blood B cells (RA from our data and a published healthy donor (31)), NR4A B cells were very rare at 0.7% −1.5% abundance. In contrast NR4A B cells were highly abundant (>40% abundance) in synovial tissue from AMP phase I (19), similar to samples in our current study (**Fig. 2E**). In the AMP phase I bulk RNA sequencing cohort (19), we found that several markers including *CD83* and *NR4A1,* which defined the NR4A^+^ cluster, were upregulated in B cells from RA biopsy samples compared to B cells from RA and OA arthroplasty samples (Fig. 2F). This NR4A cluster enrichment appeared to be driven by the inflammatory state of the tissue (1.7-fold and 2.1-fold increase in *CD83* and *NR4A1* gene expression leucocyte-rich RA vs OA, p<0.05; NS for leucocyte-poor RA vs OA) (19).

### Evidence of somatic hypermutation in synovial B cells

To define the Ig repertoire in single B cells, we performed V(D)J sequencing on the same sorted CD19^+^ cells used to generate the 5’-based gene expression library, allowing a joint analysis of transcriptional profile and immune repertoire in the same cell (Fig. 1, (fig. S2). We calculated somatic hypermutation (SHM) rates in each B cell as the number of substitutions from germline in heavy and light chains vs the total length recovered chains.

Similar to previously published data all three naïve clusters exhibited very low SHM frequencies, while memory and plasma cell clusters showed the highest mutation rates (32–34) (Fig. 3A and fig. S2F). The LMNA^+^ cluster has a mutation rate comparable to that of memory B cells, supporting that LMNA^+^ are memory B cells that may have developed in the synovium. Though mutated cells are apparent in the NR4A^+^ cluster, the average SHM rate was not significantly higher than naive B cells (1.5%, 95% CI −1%--4%) while memory and plasma cells had clear differences compared to naive, suggesting that NR4A^+^ cells may be in an earlier stage of the affinity selection and maturation process (Fig. 3A). Furthermore, we detected *IGHA* expression in a small number of NR4A B cells, providing evidence of class-switch recombination (fig. S2E).

**Fig 3.**
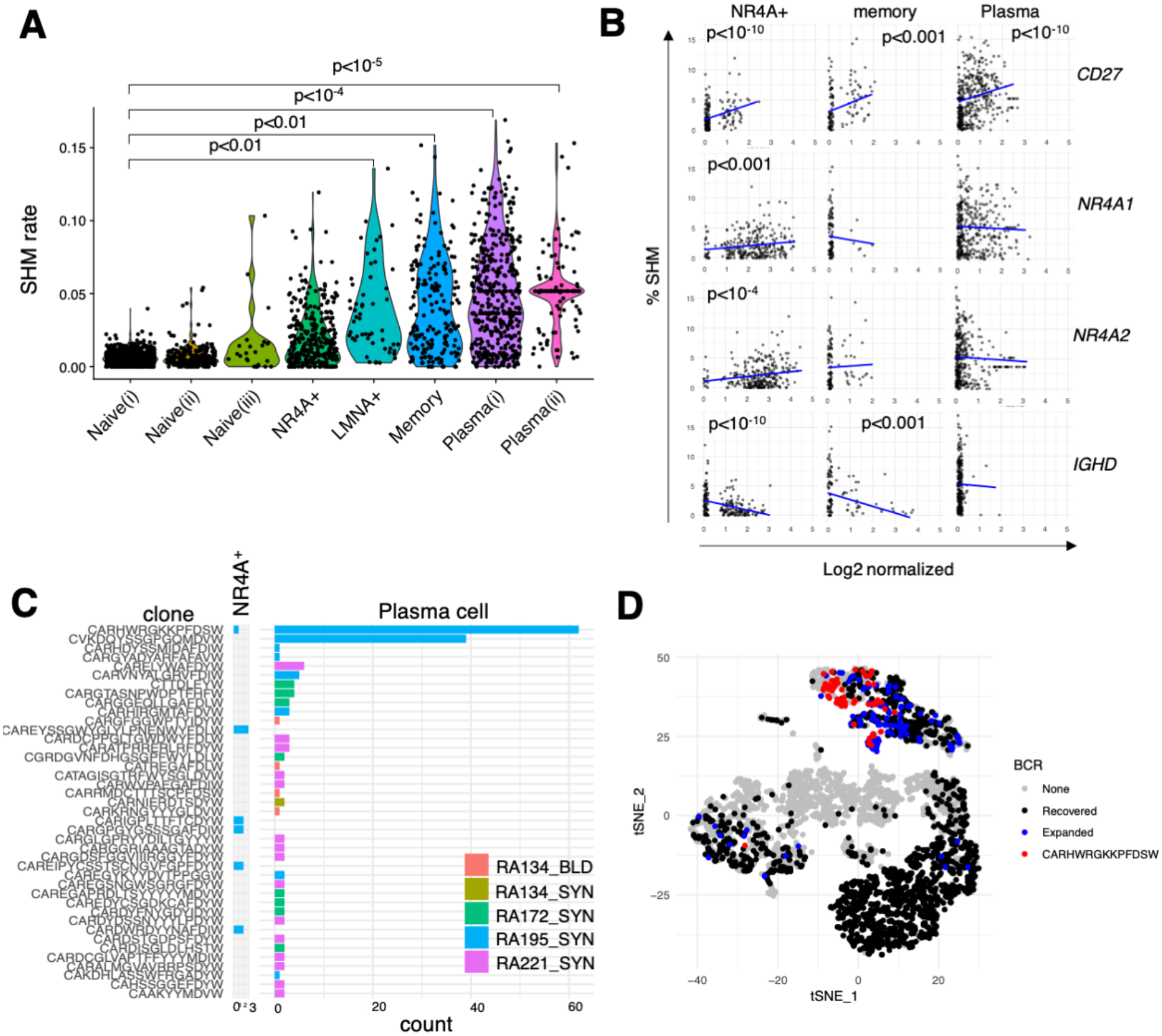
B cells in NR4A+ cluster display somatic hypermutation (SHM), clonal expansion, and shared clonality with plasma cells. (A) Violin plots display SHM rate, averaged across detected chains, in each B cell cluster obtained from single cell BCR sequencing. Each cluster’s SHM rate was tested against Naive(i) using linear mixed models, with p<0.05 displayed. (B) Scatter plots display the associations between % SHM and expression level of *CD27*, *NR4A1*, *NR4A2* and *IGHD* in NR4A^+^, memory and plasma cell (combined PC(i) and PC(ii) clusters) clusters. Blue lines display a linear mixed model fit. p-values are from a linear mixed effect model that adjusts for sample and number of genes detected. Only p-values < 0.05 are displayed. (**C**) Number of cells with >97% DNA identity of the heavy-chain CDR3. The amino acid sequence of each putative clone is shown on the left. (**D**) Heavy-chain BCR data were superimposed on cluster t-SNE plot. None (gray) indicates cells that BCR was not recovered, Recovered (black) marks cells that BCR was recovered, Expanded (blue) show cells with clonality in ≥2 cells, CARHWRGKKPFDSW (red) marks cells with a shared clone recovered from both NR4A+ and Plasma cells.

Next, we looked at the relationship between SHM rate and the expression levels of *CD27*, *NR4A* genes and *IGHD*. As expected, all subpopulations showed increased SHM with increased expression of CD27, while NR4A and memory subpopulations showed inverse associations of SHM with *IGHD*. However, particular to only the NR4A^+^ cluster, an accumulation of SHM positively associated with the expression levels of *NR4A1* and *NR4A2* (Fig. 3B), suggesting a potential role for *NR4A* genes in the RA synovial immune reaction. There was no significant association found between the expression of either the *NR4A* genes nor *IGHD* in naïve and LMNA^+^ clusters. Importantly, we detected expansion of 43 families of putative clones in the synovium and five in the peripheral blood, most prominently featured in the PC and NR4A clusters. Each clonal family was confined to a single transcriptomic subpopulation (naïve, memory, LMNA, NR4A, or PC), and a finding not be expected to occur by random chance (p<0.001 by permutation test). The only exception was the clonal family CARHWRGKKPFDSW which was detected in both the NR4A and PC cluster, consistent with a developmental relationship between these subsets (Fig. 3, C and D and fig. S2D).

### Enrichment of an activation and GC formation transcriptomic profile in the NR4A subset

To further clarify the 8 synovial B cell clusters, we performed gene set enrichment analysis (GSEA). All three naïve clusters were significantly enriched with naïve B cells from blood and tonsil (Bm1+Bm2) (Fig. 4A). In contrast, memory and LMNA^+^ clusters were enriched for gene sets expressed by tonsil Bm5 (a memory subset) and to a lesser extent memory B cells from blood, supporting their identity as memory B cells (Fig. 4A). The analysis also confirmed the identity of PC clusters (35) (Fig. 4A). The NR4A cluster was strongly enriched for genes expressed by GC LZ B cells (Fig. 4, A and C), including *EGR1/2/3, CD83, BCL2A1* and *GPR183* (36) (Fig. 4B).

**Fig 4.**
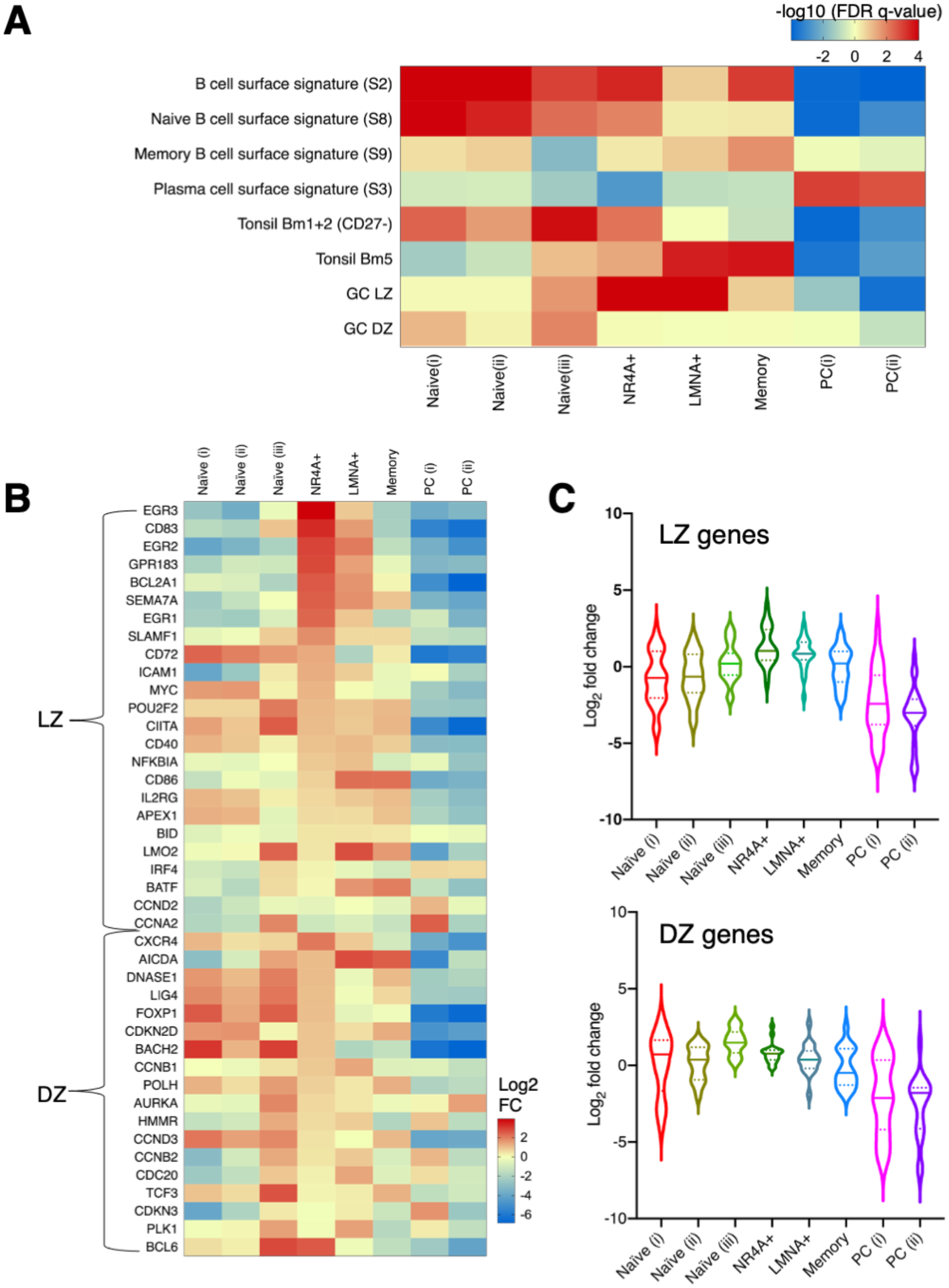
NR4A+ cluster is enriched in genes expressed by germinal center light zone (GC LZ) B cells. (**A**) Heatmap displays signed-log10 (FDR q-value) of gene set enrichment analysis between our B cell clusters and published blood and tonsil B cell subsets. (**B**) Log2 fold change of germinal center dark zone (GC DZ) and light zone (GC LZ) genes from a published study (36) in each B cell cluster. (**C**) Violin plots display log2 fold change of all LZ genes combined (top) or all DZ genes combined (bottom) from (B) in each cluster. Solid lines represent median and the dotted lines depict quartiles.

Recently, single cell RNA sequencing of GC B cells from conventional SLO has revealed a dynamic spectrum of DZ, LZ and intermediate zone B cells (37, 38). In particular, between the DZ and LZ there was a gradient of down-regulation of classic DZ genes like *CXCR4* and up-regulation of classic LZ genes like *CD83*. Along this gradient, there was an initial up-regulation of BCR signaling followed by NFkB signaling. The later stages of the intermediate zone GC B cells were characterized by up-regulation of CD40 and MYC signaling, indicative of their potential to interact with T cells (38). In our data, the NR4A B cells in RA synovial tissue displayed characteristics compatible with these intermediate cell stages. First, in contrast to classic LZ, NR4A cluster cells expressed *CXCR4*, *BCL6* and *FOXP1*, genes associated with GC DZ (36, 38, 39) (Fig. 4, B and C and fig. S3). Second, NR4A B cells showed evidence of BCR stimulation (fig. S3) and high levels of immediate early genes associated with recent BCR stimulation including *NR4A1, NR4A3, EGR1* and *c-Fos* (40–42) and NFkB signaling (fig. S3). In contrast to classic LZ cells, the NR4A cluster does not show consistent up-regulation of CD40 and MYC signaling with the exception of *BCL2A1* (CD40 signaling) and *GPR183* (MYC signaling) (fig. S3). Our NR4A B cell cluster shares remarkable transcriptomic similarity with a recently described activated B cell state in the human tonsil that appears to be on a trajectory to GC formation (34).

### Highly expressed chemokines and cytokines in NR4A B cells

Chemokines and their interaction with corresponding chemokine receptors are vital for the organization of lymphoid organs and B cell functions (15). A number of ELS promoting factors are up-regulated in lymphoid-like RA synovial tissue (43, 44). To investigate the relevant chemokine and cytokine pathways in synovial tissue, we next examined differential gene expression of chemokines, cytokines and their receptors in each B cell cluster. Both naïve and NR4A B cells showed significant enrichment for *CXCR5*, *CCR7* and *CXCR4* (Fig. 5, A and B). These chemokine receptors are instrumental to orchestrate homing to B cell follicles (*CXCR5*), migration across high endothelial venules (HEVs) (*CCR7*), and dynamic mobilization inside GC (*CXCR4*) (45, 46), suggesting that these B cells can be recruited to the synovium and localize in response to specific chemokine signals. Receptors for B cell survival factors are also expressed by multiple B cell subsets. Naïve and NR4A subsets showed significant up-regulation of *TNFRSF13C* (BAFFR), while LMNA^+^ and memory shared up-regulation of *TNFRSF13B* (TACI), a receptor that can bind to both B cell survival factors BAFF and APRIL. Consistent with previous reports, PC uniquely showed up-regulation of *TNFRSF17* (BCMA), a BAFF receptor important for PC survival (Fig. 5A). Furthermore, NRA4^+^, LMNA^+^ and memory showed significant enrichment of *TNFRSF1* (TNFR2), a receptor for TNF (Fig. 5A). We also observed that LMNA^+^ and memory subsets uniquely up-regulate *SIGIRR*, a negative regulator of IL-1 receptor signaling pathway implying response to IL1 family cytokines (Fig. 5A).

**Fig 5.**
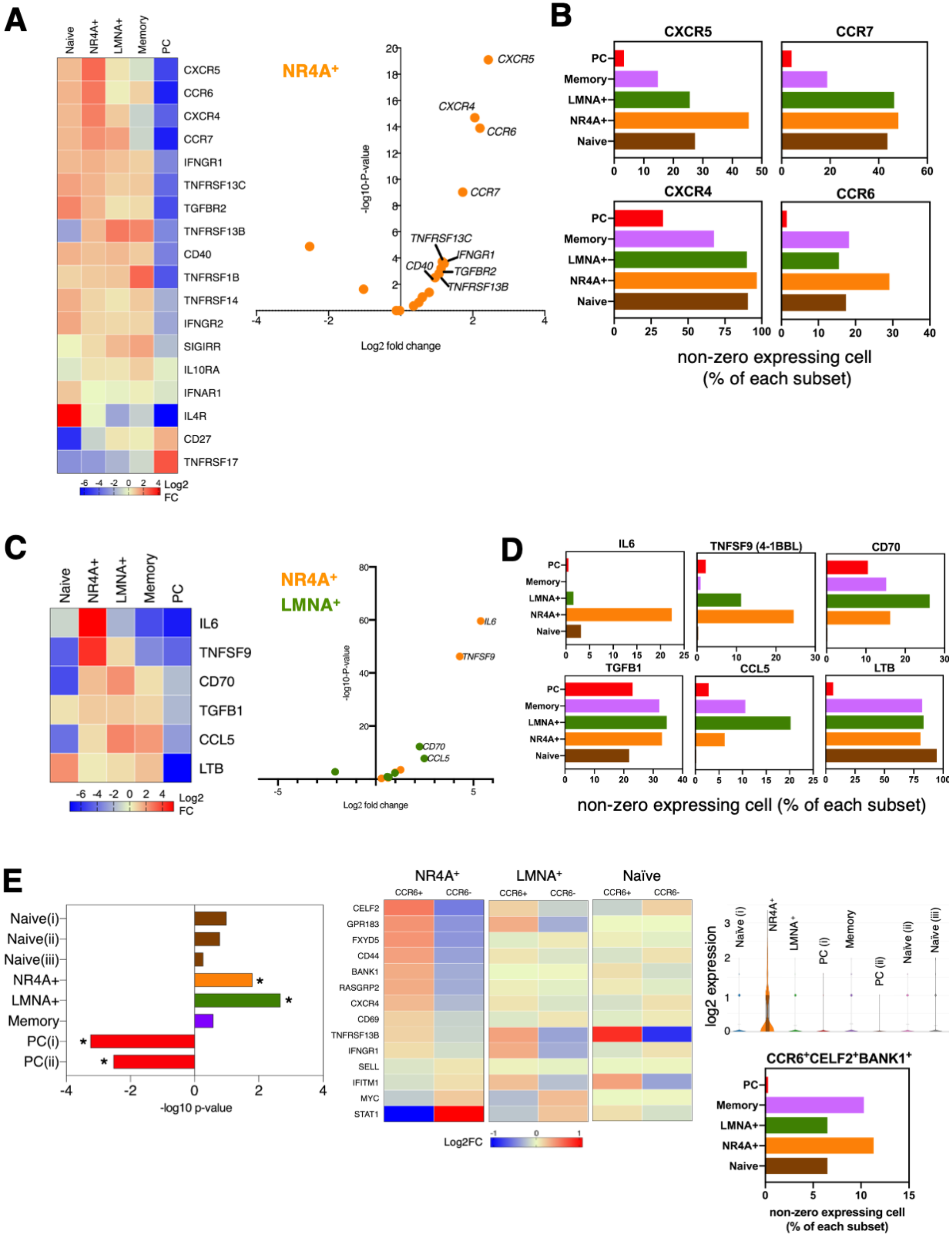
NR4A+ cluster expresses chemokine receptors and cytokines important for ELS formation. (**A**) Heatmap displays log2 fold change of cytokine and chemokine receptors in each B cell subset (n=1,250 naive, 746 NR4A^+^, 633 LMNA^+^, 330 memory and 779 PC). Volcano plot display log2 fold change and –log10 p value of genes in the heatmap. (**B**) Percentage of cells of each subset that express *CXCR5, CCR7, CXCR4* or *CCR6*. (**C**) Heatmap displays log2 fold change of cytokines and chemokines in each B cell subset. Volcano plot display log2 fold change and –log10 p value of genes in the heatmap in NR4A^+^ and LMNA^+^ subset. For A and C, shown are genes that are expressed in at least 20% of the cells of at least one subset. p-values are adjusted using the Benjamini-Hochberg correction for multiple tests. (**D**) Percentage of cells of each subset that express *IL6, TNFSF9, CD70, TGFB1, CCL5* or *LTB*. (**E**) GSEA analysis revealed significant enrichment of PreM signature in NR4A^+^ and LMNA^+^ subsets,*p < 0.05. Heatmaps show log2 fold change of PreM genes (38) in CCR6^+^ and CCR6^-^ cells within NR4A^+^, LMNA^+^ or naïve subset. Violin plots showed log2 expression of CCR6 in each cluster. Bottom right bar graph shows percentage of cells of each subset that co-express *CCR6, CELF2* and *BANK1.*

B cells can play a direct role in the organization of SLO by producing lymphotoxin *β* (47). More recently, B cell-derived IL6 has been implicated in spontaneous GC formation in systemic lupus (48). We found that all non-PC B cell subsets express lymphotoxin *β*(Fig. 5, C and D). However, the NR4A^+^ subset uniquely expressed *IL6*, and co-expressed *LTB* (Fig. 5C). As previously noted, *GPR183* (EBI2) is highly expressed in the NR4A subset. GPR183 has pleiotropic roles in B cell positioning depending on the balance of other chemokines. *GPR183* together with *CXCR4*, also expressed at high levels by the NR4A subset, are critical factors for the segregation of GC B cells in the DZ and LZ (Fig. 5A). NR4A B cells also have elevated expression of a number of co-stimulatory and signaling molecules important for putative cell-cell interactions with APCs and T cells. In addition to *IL6* (interaction with IL6R on APCs and Tfh), this includes high levels of *TNFSF9* (4-1BBL) (Fig. 5, C and D), a ligand that binds to 4-1BB on activated CD4 and CD8 T cells to provide co-stimulatory signals (49, 50). Overall, these data suggest NR4A B cells may be a critical initiator of ELS formation in synovial tissue. Independent analysis of the RA cohort from AMP also supports these findings (fig. S5). Of note, we observed the presence of histologic ELS in our samples by immunofluorescent staining with classic GC markers (fig. S4).

One of the differentially expressed chemokine receptors in the NR4A subset is CCR6 (Fig. 5, A and B). Single cell analysis of GC B cells from human tonsil has identified a distinct cell cluster of memory B cell precursors (PreM), originating within the LZ that shares a gene signature with previously reported murine CCR6^+^ PreM B cells (38, 51). The range of *CCR6* expression within the NR4A^+^ cluster (Fig. 5E) suggests that memory precursor cells are originating here and is consistent with memory precursor commitment to post-GC differentiation in the early stage of the LZ (38). Consistent with *CCR6* expression, we found that NR4A^+^ and LMNA^+^ cells are significantly enriched in a PreM signature (Fig. 5E). In contrast to naïve subsets, CCR6^+^ cells in the NR4A^+^ subset showed higher expression of most of the genes described as a signature of PreM including *CELF2*, *BANK1* and *CD44* compared to CCR6^-^ counterparts, while LMNA^+^ CCR6^+^ displayed a smaller subset of PreM signature genes (Fig. 5E). In addition, the NR4A subset contained high fractions of cells co-expressing *CCR6, BANK1* and *CELF2* (Fig. 5E).

### A gene expression activation continuum from naïve to NR4A^+^ B cell

The variability in *NR4A1, NR4A2, IGHD* and *CD27* expression in the NR4A subset (Fig. 2, B and D) led us to hypothesize that B cells in synovium might exist along a continuum from a naïve state. Projecting the gene expression data of the B cells onto two dimensions using principal component analysis, we sought to further define the relationship between B cell subsets. Plasma cells were excluded from this analysis. Dimension 1 (4% variance) separated NR4A cells from naïve. In contrast, dimension 2 explained the difference between the NR4A and memory subsets (2% variance) (fig. S6A). LMNA^+^ cells were positive in both dimension 1 and 2. In dimension 1, we found that genes with large gene loadings are the ones involved in lymphocyte activation and GC molecular events including *CDKN1A*, *NFKBID*, *GPR183*, *CD83*, *LY9*, *KLF6* and *IGHG1*, and are expressed in increasing levels from naïve to NR4A^+^ and LMNA^+^ clusters. Consistently, the NR4A cluster is significantly enriched for gene sets associated with cell activation (fig. S6D). Additionally, a number of DNA-binding transcription factors (*NR4A2, NFKBID, NR4A3, JUND, DNAJA1, CREM, FOS*) gradually increase from naïve to NR4A cluster, while genes characteristic of naïve B cells (*TCL1A, IGHM, FCER2*) are down-regulated (fig. S6B). We observed along dimension 1 cells that are in transition states between naïve and NR4A^+^. Thus, expression of NR4A^+^ markers including *NR4A1* and *DUSP1* are gradually increased, while concurrently markers associated with naïve status such as *TXNIP* and *CD79B* showed a gradual decrease in expression (fig. S6B). In dimension 2, genes with increased expression levels are those associated with memory B cells including *HOPX*, *IGHA1*, *S100A4* and *CD99*. We observed evidence of transitioning between NR4A^+^ and memory state with increased expression of memory genes (*S100A6*, *HOPX*) and decreased expression of NR4A^+^ genes (*JUN*, *DUSP1*). Our data are consistent with a continuum of states from naïve to NR4A^+^ with both loss of naïve and acquisition of activation status. Whether LMNA^+^ cells differentiate from memory, or NR4A^+^ or both remains to be defined.

### NR4A1 protein expression in SLO and synovial ELS

Endogenous NR4A1 (Nur77) expression is an indicator of antigen receptor signaling in both mouse and human B cells (52–54). We first defined endogenous NR4A1 expression in classic human SLO. We detected NR4A1^+^ B cells in GCs in close proximity to CD21^+^ LZ follicular dendritic cell networks (FDC) inside GC (Fig. 6A). Interestingly, we also observed NR4A1 expression in CD138^+^ plasma cells surrounding GCs (Fig. 6B). Additional analysis by flow cytometry of tonsil B cells shows that multiple subsets (Bm1+Bm2, Bm5, GC and PC) of tonsil B cells express NR4A1, but with significantly more NR4A1^+^ cells in the GC subset (Fig. 6C). Paralleling immunofluorescent staining, we identified NR4A1^+^ PCs by flow cytometry (Fig. 6C). Furthermore, when we subset GC B cells based on CD83 and CXCR4, significantly more NR4A1^+^ cells were present in CD83^+^CXCR4^+^ intermediate GC B cells than in CD83^-^CXCR4^+^ (DZ), CD83^+^CXCR4^-^ (LZ) or CD83^-^CXCR4^-^ GC B cells, consistent with our RNA sequencing data that synovial NR4A^+^ B cells express high levels of both *CXCR4* and *CD83* (Fig. 6D).

**Fig 6.**
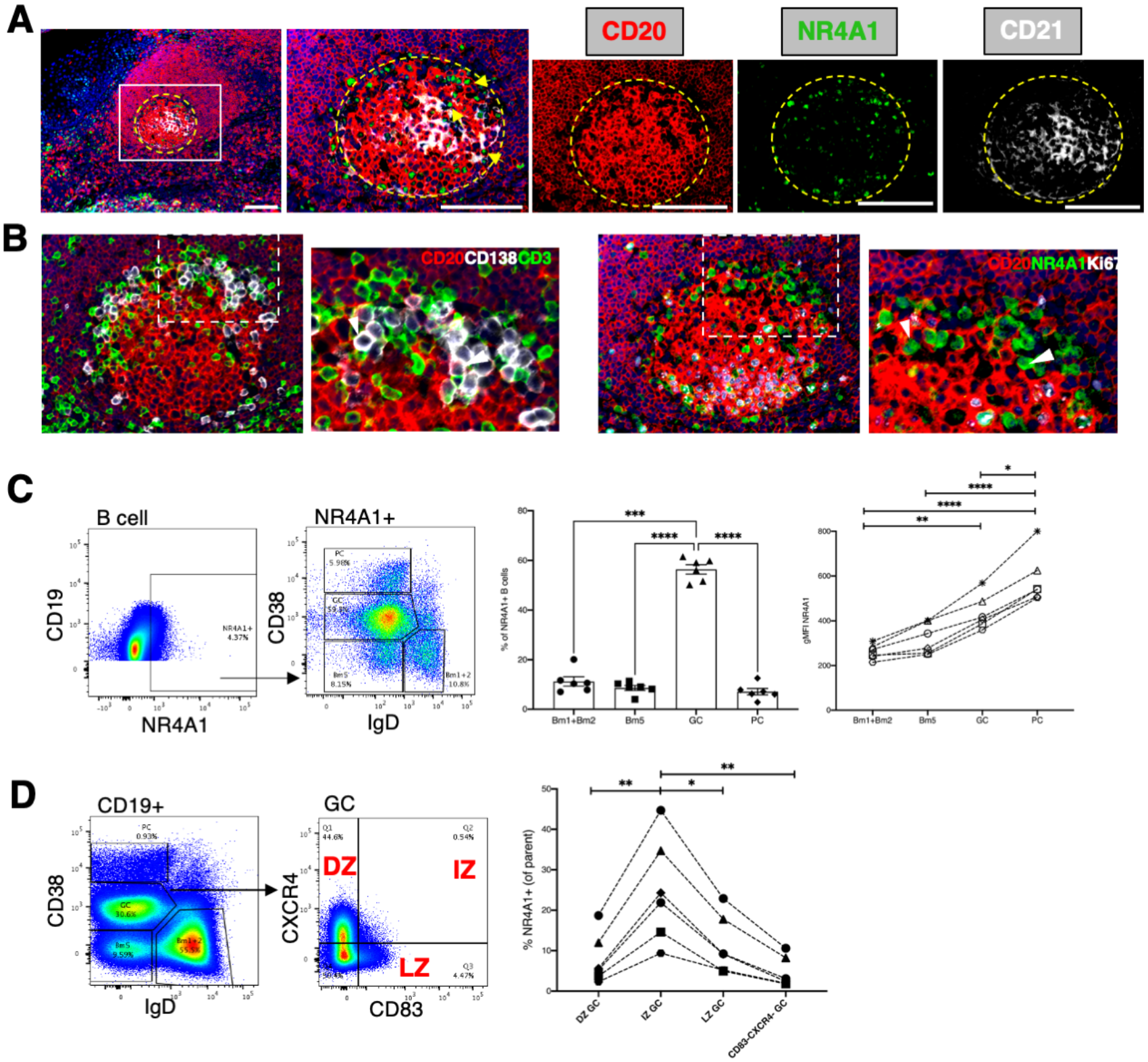
NR4A1 is expressed by B cells and plasma cells in tonsil SLO. (**A**) Immunofluorescent (IF) staining of NR4A1 (green), CD20 (red) and CD21(white) in tonsil. Representative 3×3 200X mosaic (left) and 200X magnification show the composite and individual channels. Box area was magnified. Dashed yellow lines outline the tonsillar GC. Scale bars = 100 μm. Yellow arrows show NR4A1^+^ B cells next to CD21^+^ FDC in the GC LZ. (**B**) IF staining of two tonsil serial sections, CD20+CD138+CD3 (left images) and CD20+NR4A1+Ki67 (right images). White arrows point out CD138^+^ plasma cells in the first section and the corresponding NR4A1^+^ plasma cells in the adjacent section. (**C**) Flow cytometry for NR4A1 expression in tonsil B cells (n=6). Dot plot representative of phenotype of NR4A1^+^ tonsil B cells with respect to CD38 and IgD expression with the percentage display in bar graph. Line graph shows geometric mean fluorescent intensity (gMFI) of NR4A1^+^ Bm1+Bm2, Bm5, GC or PC. (**D**) GC B cells were further characterized based on CXCR4 and CD83 expression. Percentage of NR4A1^+^ cells were determined in each CD83-CXCR4 population and plotted per sample. For C and D line graphs, dash line connect data from the same sample. Error bars are SEM. * p < 0.05, ** p < 0.01, *** p < 0.005, **** p < 0.0001 by non-parametric one-way ANOVA and Tukey’s multiple comparisons test.

To investigate endogenous NR4A1 protein expression in synovial B cells, we performed immunofluorescent staining on synovial tissue sections. Interestingly, the most intense NR4A1 expression in the synovium is in PCs (Fig. 7A), consistent with the tonsil staining and NR4A1 expression in a PC subpopulation in the single cell RNA sequencing data (Fig. 2B). We also observed NR4A1^+^ B cells in aggregates (Fig. 7A), lacking Ki67 expression, in agreement with a GC intermediate zone. Flow cytometry of B cells from disaggregated RA tissue and synovial fluid also revealed high NR4A1 expression. Based on surface markers, 8.5% of NR4A1^+^ cells in synovium are IgD^-^CD27^++^, 42% are IgD^-^CD27^+^, 36% are IgD^-^CD27^-^, 6.3% are IgD^+^CD27^+^ and 7.2% are IgD^+^CD27^-^ (naïve) B cells (n=7, Fig. 7, B and C). In marked contrast, only a small percentage of peripheral blood B cells (from healthy controls or RA) spontaneously express NR4A1 (Fig. 7C).

**Fig 7.**
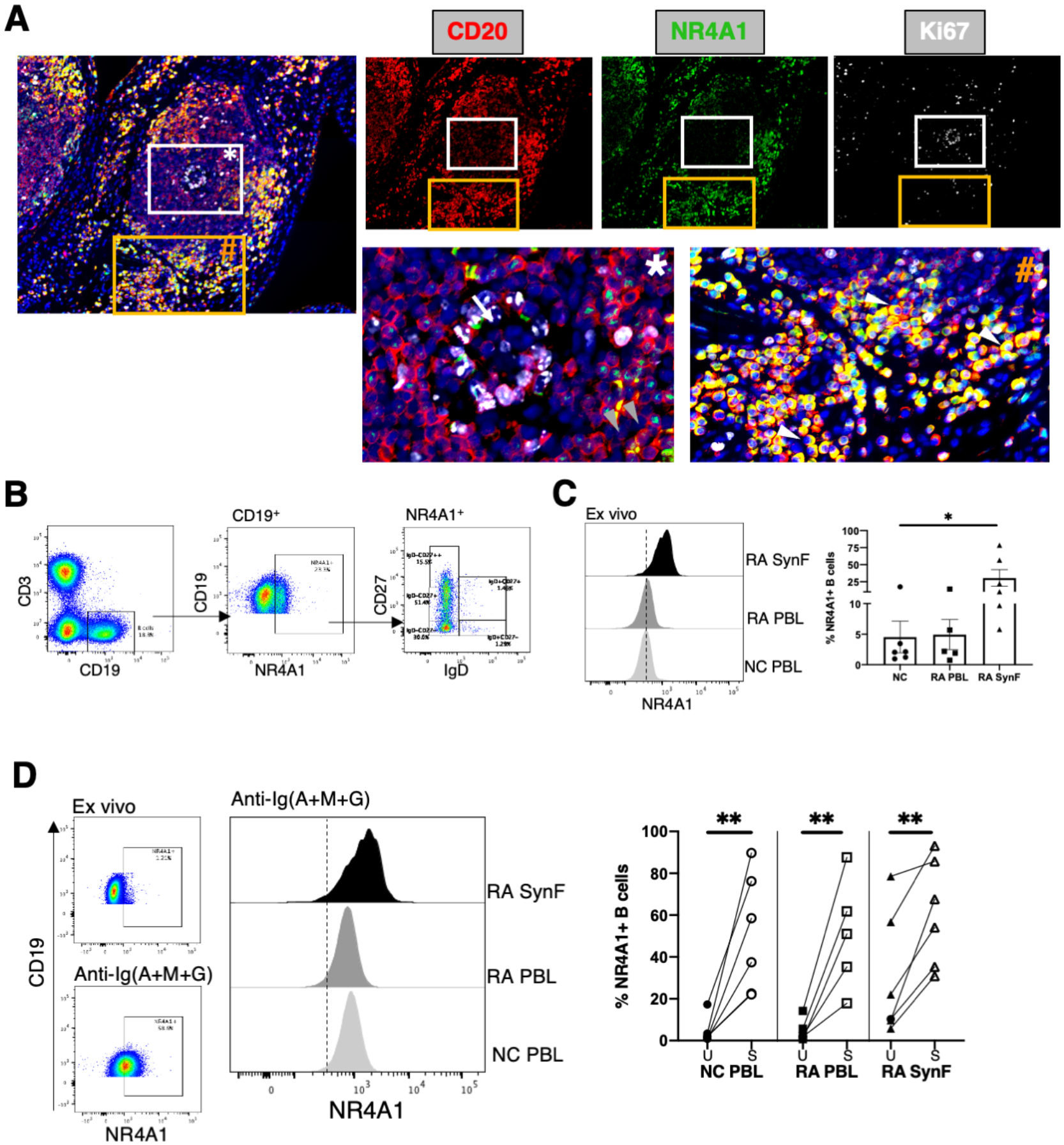
NR4A1 is expressed by B cells and plasma cells in synovium ELS. (**A**) IF staining of NR4A1(green), CD20 (red) and Ki67 (white) in synovial serial sections. Representative 3×3 200X mosaic show the composite and individual channels. White (*) and orange (#) boxes are magnified. White arrow points out an aggregate of Ki67^+^ B cells in an ectopic GC DZ. Gray arrows show NR4A1^+^ B cells outside of GC. White arrow heads on the right most image show NR4A1^+^ plasma cells. (**B**) Example dot plot showing spontaneous NR4A1 expression by synovial B cells and plasma cells. (**C**) Histograms of ex vivo NR4A1 expression from RA and normal control (NC) peripheral blood (PBL) B cells and RA synovial fluid B cells (RA SynF). Dash line is the median MFI of NR4A1 in NC PBL. Bar graph represents mean of % NR4A1^+^ B cells in ex vivo sample from NC (n=5), RA (n=5) and synovial fluid (n=6). Error bars are SEM and * p < 0.05 by non-parametric one-way ANOVA and Tukey’s multiple comparisons test. (**D**) Histogram represents NR4A1 expression in B cells from PBMC treated with anti-Ig(A+M+G) for 4 hours. Dash line is isotype control for NR4A1. Scatter plots represent % NR4A1^+^ B cells after treatment (S) compared to untreated control (U). ** p < 0.01 by paired t-test.

When we stimulated peripheral blood and synovial B cells with a cocktail of anti-IgG, anti-IgA and anti-IgM to engage BCR of multiple isotypes, NR4A1 expression was significantly upregulated (Fig. 7D). The NR4A1 mean fluorescent intensity (MFI) was significantly increased in stimulated synovial B cells compared to untreated and stimulated peripheral blood healthy and RA B cells (Fig. 7D). In addition to NR4A1 up-regulation, we also found that BCR stimulation up-regulated NR4A2 and NR4A3 expression at the mRNA level (fig. S7).

### Lymphoid specific NR4A subset genes correlate with RA synovial tissue pathotype and increase in the blood during RA flare

Next, we sought to relate NR4A subset signature genes to synovial histology and clinical characteristics in an independent cohort of synovial biopsies from treatment-naive patients: The Pathobiology of Early Arthritis Cohort (PEAC) (9). Based on histology scores, synovial samples were classified as lympho-myeloid (CD20 B cell aggregate rich), diffuse-myeloid (CD68 rich in the lining or sub-lining layer but poor in B cells), or fibroid (paucity of immune-inflammatory cell infiltrate), as previously described (9). We focused on the expression of *CD83* and *GPR183* as lymphoid specific genes representative of the NR4A cluster as RNA sequencing of total tissue is available in this study (Fig. 8, A and B). Both genes were strongly enriched in the lymphoid pathotype tissues and also showed significant correlation with synovial histology scores for CD20, CD3, CD138, and sub-lining CD68. These data suggest that infiltration of multiple immune cell types associated with ectopic lymphoid responses in the synovial tissue may be linked to the B cell NR4A subset. Additionally, *CD83* expression showed a relationship with ultrasonographic joint synovial thickness confirming that ELS gene expression strongly matches imaging signs of active joint inflammation in the particular joint undergoing biopsy. Surprisingly, *NR4A1* and *NR4A2* whole synovial gene expression was lower in the lymphoid vs fibroid pathotype (p=0.00025 for *NR4A1* and p=0.22 for *NR4A2*) and inversely correlated with histologic immune cell infiltration (*NR4A1*: p=0.00051(CD3), p=1.7×10^-5^(CD20), p=2.3×10^-5^(CD138); *NR4A2*: p=7.5×10^-5^(CD3), p=2.6×10^-5^(CD20), p=5.2×10^-5^(CD138)) consistent with B cell independent roles for the NR4A family in the synovium in other cell populations that may be anti-inflammatory (55).

**Fig. 8.**
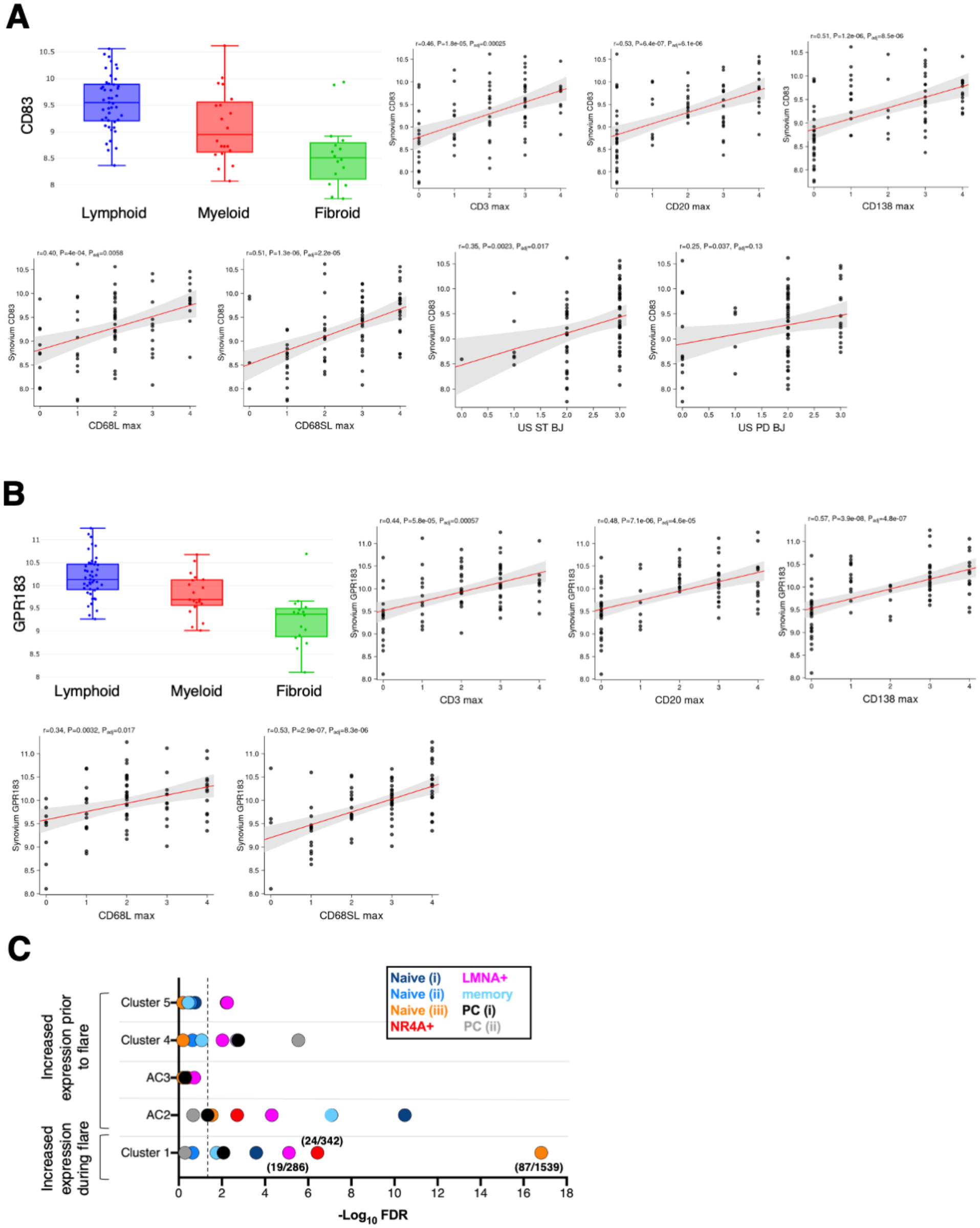
Lymphoid specific NR4A subset genes correlate with RA synovial tissue pathotype and increase in the blood during RA flare. Boxplots and linear regression of histology and clinical parameters by tertile demonstrating correlation with (**A**) CD83 (P_adj_ lymphoid v myeloid=3.0e-0.2; P_adj_ myeloid v fibroid=3.4e-0.2; P_adj_ lymphoid v fibroid=3.3e-0.7) and (**B**) GPR183 (P_adj_ lymphoid v myeloid=5.4e-0.3; P_adj_ myeloid v fibroid=4.6e-0.2; P_adj_ lymphoid v fibroid=4.1e-0.8). Ultrasound biopsy joint parameters: (ST, synovial thickness; PD, power doppler) at the biopsy joint (Ultrasound ST/PD BJ). p values were calculated by linear regression models. The plots and p-values were obtained from https://peac.hpc.qmul.ac.uk/ (9). (**C**) The plot shows significance of enrichment scores of marker genes from various synovial B cells in clusters of genes differentially expressed over time to flare (20). The dash vertical line represents the threshold for significance (-log_10_ FDR > 1.3). The numbers in parentheses display the gene ratio (# overlapping genes/total genes in the set) of LMNA^+^, NR4A^+^ and Naïve (iii) with Cluster 1.

Although our initial data suggested that NR4A B cells are rare in RA blood (**Fig. 7C**), we wondered whether circulating numbers of this activated B cell population may vary over time depending on disease activity. To begin to understand this dynamic, we compared our synovial B cell cluster signatures with blood transcriptional profiles recently defined relative to flare (20). In this data set two transcriptomic clusters appeared in the blood preceding flare: AC2 was increased 2 weeks before flare and AC3 was increased 1 week before flare. AC2 has characteristics of naïve B cells, while AC3 is a unique CD45−CD31−PDPN+ mesenchymal population, termed PRIME (preinflammatory mesenchymal) cells (20). An additional cluster 1 was increased at the onset of flare. When we examined the overlap between these blood clusters and our single cell B cell clusters (Fig. 8C, and table S2) using hypergeometric tests for enrichment, we found that AC2 was significantly enriched with transcripts characteristic of peripheral blood dominant naïve B cell clusters, though the synovial NR4A and LMNA clusters were represented (Fig. 8C). In contrast, the flare cluster 1 was significantly enriched with naïve (iii), NR4A, and LMNA synovial populations (7%, 6.6%, and 5.6% overlapping markers with the NR4A, LMNA and Naive (iii) clusters, respectively). This is consistent with a trajectory of B cell activation whereby resting blood B cells are recruited from the blood to the synovium pre-flare and then activated within the synovial microenvironment to express NR4A during flare.

## DISCUSSION

Our integrated single-cell analysis of RA synovial B cells has revealed a surprising level of B cell activation, with a large fraction of synovial B cells demonstrating evidence of recent antigen stimulation as revealed by upregulation of the NR4A gene family. NR4A B cells appear primed for cell-cell interactions with APCs and T cells and, further, have evidence of molecular events associated with active GC reactions, including class-switch recombination, somatic hypermutation, and clonal expansion with selection into the PC compartment. Our findings suggest a spectrum of activation of B cells within the synovial microenvironment with a pluripotent fate that includes differentiation to memory or PC lineage *in situ*.

Prior studies have supported local synovial activation of RA B cells with evidence of SHM and clonal expansion (8, 17) using a number of approaches including enzymatic digestion or focal microdissection with bulk or single cell BCR sequencing. An important advance in the current study is the high-throughput combination of single cell RNA sequencing and Ig gene repertoire. This allowed the unprecedented identification of a spectrum of B cell activation in synovial B cells downstream of BCR signaling as revealed by *NR4A1* expression. A key finding from our study was the relationship between the degree of NR4A family gene expression and SHM. Notably, this dynamic process was unique to the synovium and not observed in B cells from RA peripheral blood or lupus kidney. NR4A1-3 encode a small family of orphan nuclear hormone receptors (NUR77, NURR1, and NOR1 respectively) which are rapidly induced by acute and chronic antigen stimulation in B cells and T cells (54, 56). Our results raise key questions regarding the precise synovial microenvironmental signals that promote NR4A family expression and 2^nd^ signals that may direct subsequent differentiation to memory B or plasma cell. Another study did combine transcriptomic and repertoire analysis of peripheral blood RA B cells and interestingly suggested that ACPA enriched B cell responses are imprinted with a T cell dependent transcriptional network (7). Given the strong association of NR4A1 expression in B cells and BCR engagement (52), we speculate that the enrichment of B cell NR4A1 expression in the RA synovium represents chronic auto-antigen stimulation and an ACPA response. This would be in keeping with other data in the literature indicating that B cells differentiating within ELS frequently target citrullinated peptides (17). However, the antigen specificity of the NR4A B cells in our study remains to be determined.

In terms of B cell fate, it is notable that NR4A family members may trigger B cell apoptosis after encounter with self-antigen in the absence of a 2^nd^ signal provided by T cell (or other) co-stimulation and act as a negative regulator of B cell activation (57). Thus, in murine B cells, the NR4A family, particularly NR4A1/NUR77, actively participates in B cell tolerance. NUR77 is up-regulated in self-reactive B cells in response to chronic antigen stimulation and selectively restricts survival by limiting B cell survival factors such as BAFF (58). In addition, a small subset of GC LZ B cells exhibiting a high SHM rate, up-regulate NR4A1/NUR77, suggesting active BCR signaling and a role for NR4A1 in selection and regulation of GC differentiation pathways into the PC compartment (53). Recently, NR4A1 and NR4A3 have been identified as partially redundant mediators of an antigen-dependent negative feedback loop in B cells to reinforce tolerance by increasing B cell dependence on T cell help and regulating clonal competition, when T cell help is limited (57). These data point to an expanding role for the NR4A family in adaptive immune response occurring in SLO. However, the role of NR4A in ELS and autoimmunity had not been studied previously. Whether NR4A attempts to restrain B cell activation within the synovium, similar to its role in a recently reported anti-inflammatory synovial tissue resident macrophage population (55) and exhausted T cells in the tumor microenvironment (59, 60), remains to be defined.

Our finding of shared clonality between the NR4A B cell and PC clusters suggests that some NR4A B cells do indeed receive the signals required to drive their differentiation to memory and PC *in situ*. The signals that control differentiation of activated B cells into memory or plasma cells in ELS require further study, but similar to SLO may depend on BCR affinity and the strength and stability of their interaction with T cells (34, 51, 61). We identified several genes within the NR4A cluster that may facilitate T-B cell interactions including CD69 (which prevents activated lymphocyte egress from SLO), CD86 (a ligand for CD28 on T cells), IL6 (which can promote both APC and T cell interactions), and ICAM1 (which can support B cell interaction with T follicular helper cells) (62). This is also in accord with a recent report showing that tetramer identified ACPA positive B cells in RA blood and synovial fluid express T cell-stimulating ligands and a persistently activated phenotype suggestive of continuous antigenic triggering (63). A key role for B cells in RA flare was also suggested by the recent discovery of a circulating expanded B cell subset (AC2) prior to emergence of a circulating CD45-/CD31-/PDPN+, PRe-Inflammatory MEsenchymal (“PRIME”) cell in RA patient blood (20). We find that this pre-PRIME circulating B cell cluster (AC2) was enriched with resting naïve (naïve (i), naïve (ii)) and memory B cells that were dominant in the blood sample we included in our single cell analysis. In contrast, the blood transcriptomic cluster appearing during flare (cluster 1) was enriched in the NR4A, LMNA, and naïve (iii) clusters. This is consistent with a trajectory of B cell activation whereby blood B cells are recruited to the synovium pre-flare, activated within the synovial microenvironment to express NR4A, and then possibly re-circulate during flare. Re-circulation of B cells between the synovium and the periphery is also supported by a recent publication demonstrating shared clonal families between these compartments in RA (64). The origin of the B cells increasing in the blood pre-flare remains to be determined.

B cells may contribute to RA pathogenesis in multiple ways, not only as the precursors of PCs and production of autoantibodies, but also by antibody independent functions including antigen presentation and cytokine production. B cells can produce cytokines and chemokines that promote ELS. Thus, lymphotoxin *β*expression by B cells has been shown to support ELS formation in a mouse colitis model (18). Lymphotoxin *β* and *α* are elevated in RA synovium and relate to the level of inflammation in the tissue (65), and LT-*β* was expressed by synovial B cells but also other cell populations (11). The precise role of B cells in ELS formation in RA synovial tissue and which sub-populations may be critical is unclear. We found a significant relationship between NR4A B cell expression and the degree of B cell infiltration and lymphoid organization in tissue implicating NR4A B cells as a potential ELS initiating and promoting B cell subset. In accord with this, NR4A B cells expressed LT-*β* and were the dominant source of IL6, the latter important for Tfh interactions and demonstrated to promote spontaneous GC formation in SLE (48).

Overall, our results highlight the unique enrichment of NR4A expressing B cells in the RA synovium and their identity as an antigen experienced B cell in the synovial ELS with the capability of differentiating to plasma cell or memory B cell in situ depending on additional signals. NR4A may serve as a biomarker of chronic autoantigen stimulation in the synovial microenvironment and a novel therapeutic target.

## MATERIALS AND METHODS

### Patient samples

Arthroplasty samples were acquired from RA patients after surgical removal as part of standard of care at Hospital of Special Surgery, NY (n=3) or collected as part of the AMP network (ImmPort SDY998) (n=1). Peripheral blood was collected prior to the procedure. Written informed consent for research use of patient samples was obtained prior to study inclusion at the time of sample collection. The study received institutional review board (IRB) approval at each site. Synovial tissue fragments were cryopreserved in CryoStor CS10 (BioLife Solution) for subsequent disaggregation at University of Rochester (66). PBMC were isolated from whole blood by Ficoll-hypaque density gradient centrifugation and cryopreserved in CryoStor CS10 (BioLife Solution). Samples from the AMP network were collected as part of multi-center cross-sectional study of patients undergoing elective surgical procedures and a prospective observational study of synovial biopsy specimen from patients with RA aged ≥ 18 years, with at least one inflamed joint (19). This cohort included synovial tissues from 16 RA biopsies, 13 RA arthroplasties and 10 OA arthroplasties. 14 RA biopsies have B cell single cell RNA sequencing data available for analysis. The Pathobiology of Early Arthritis cohort (PEAC) contains data from 90 RA patients fulfilling 2010 ACR/EULAR RA Classification Criteria that were enrolled as part of the Medical Research Council funded multi-center PEAC project (9) (https://peac.hpc.qmul.ac.uk/). Normal control and RA peripheral blood were obtained from patients at University of Rochester under IRB approved study protocols. PBMCs were isolated by Ficoll-hypaque density gradient centrifugation and cryopreserved in freezing medium (90%FBS/10%DMSO) until used. Tonsil tissue samples were obtained as discarded tissue from tonsillectomy procedures. Portions of tissue was fixed in formalin and paraffin-embedded for histologic staining. The remaining tissue was mechanically disaggregated to obtain cell suspension. The cell suspension was further purified by Ficoll-hypaque density gradient centrifugation. Isolated mononuclear tonsil cells were cryopreserved in freezing medium (90%FBS/10%DMSO) until used.

### Preparation of synovial tissue and PBMC and flow cytometry cell sorting

Cryopreserved synovial fragments were thawed per SOP (66). Synovial tissue fragments were then disaggregated by combination of an enzymatic digestion and mechanical disruption as described. PBMCs were thawed by rapidly warming the cryovial in a 37 °C water bath. To stain cells for sorting, cells were first incubated with Fc-Block (Biolegend) for 5 minutes at room temperature (RT). Cells were then incubated with a mixture of fluorochrome-conjugated antibodies (table S3) for 30 minutes on ice, washed with PBS and then resuspended in PBS solution containing DAPI (Biolegend). Cells were sorted on a FACSAria II (BD biosciences) into 5%FBS/RPMI media using a 100-μm nozzle. B cells were gated as CD45^+^CD3^-^CD14^-^CD19^+^ (fig. S1A).

### Single cell RNA and BCR repertoire sequencing

Cellular suspensions were loaded on a Chromium Single-Cell Instrument (10x Genomics, Pleasanton, CA, USA). Single-cell RNA-Seq libraries were prepared using Chromium Single-Cell 5′ Library & Gel Bead Kit (PN-1000006, PN-1000014, 10x Genomics). The barcoded, full-length V(D)J segments were enriched by PCR with primers specific to the BCR constant regions using the Chromium Single Cell V(D)J Enrichment Kit, Human B cell (PN-1000016, 10x Genomics). For both B-cell enriched library and 5’ gene expression library, enzymatic fragmentation and size selection was used to optimize the cDNA amplicon size and indexed sequencing libraries were constructed by End Repair, A-tailing, Adaptor Ligation, and PCR. Final libraries contain the P5 and P7 priming sites used in Illumina bridge amplification. Paired end reads of 150nt were generated for each B-cell enriched library using the MiSeq sequencer (Illumina). The 5’ gene expression libraries were sequenced following 10X Genomics read length guidelines on the NovaSeq6000 sequencer (Illumina).

### Single cell RNA and repertoire sequencing analysis

Raw reads generated from the Illumina basecalls were demultiplexed using Cell Ranger mkfastq version 2.1.1 utilizing bcl2fastq version 2.19.1.403. Cells were aligned and counted against the Cell Ranger reference refdata-cellranger-GRCh38-1.2.0 using Cell Ranger count version 2.1.1, utilizing only the library index, project, and sample ID for a given sample. We performed quality control to remove outlier events and doublets based on the cell’s total UMI size versus the number of distinct genes expressed in SeqGeq^TM^ v1.4 (FlowJo, LLC) by gating on bivariate scatter plots. We also filtered putatively degraded cells based on the proportion of UMIs in the cell derived from the mitochondrial genome. These steps removed 252 cells. After filtering, the data were exported for further analysis in scran 1.8.4 and Seurat 2.3.4 (22). The top 1000 variable genes, excluding IGHV, IGKV and IGLV genes were used to derive 6 principal components. These were used to define clusters using the Louvain method and resolution parameter 0.5. Based on expression of non-B cell markers (*COL1A2*^+^, *FN1*^+^, *MS4A1*^-^), we identified 77 putative fibroblasts, and accordingly excluded these cells from further analyses. BCR clonotypes were assigned by requiring 97% identity of the DNA sequence of the CDR3 region of the heavy or light chain using CellaRepertorium version 0.8.1. (http://bioconductor.org/packages/release/Bioc/html/ CellaRepertorium.html). For somatic hypermutation (SHM) rate calculation, the consensus FASTQ sequences were aligned to the human immunoglobulin reference from the international ImMunoGeneTics information system (IMGT) using HighVQuest to calculate the percent identity in V and J in heavy and light chains which then averaged, weighting by the length of each segment.

### Projection of clustering using SingleR

Gene-cell count matrices were downloaded for the AMP Phase I RA and SLE cohorts (ImmPort accessions SDY998 and SDY997). Filtered gene-cell count matrices for CD19^+^ B cells from Zheng et al 2017 (31) were downloaded from https://support.10xgenomics.com/single-cell-gene-expression/datasets/1.1.0/b_cells. 12848 gene symbols expressed in common in these datasets and our data were retained and the data merged. QC was evaluated using the quickPerCellQC function in scran and 874 were cells filtered. A SingleR model was trained on our data and its 8 cluster labels, and the inferred cluster identities in the other datasets were queried. Binomial mixed models were used to estimate the frequency of the NR4A^+^ cluster among these other data sets. For bulk RNAseq from AMP RA Cohort, data was downloaded (ImmPort accession SDY1299). DESeq2 version 1.20.0 was used to evaluate differential expression of NR4A^+^ marker genes in sorted B cells from this study.

### Gene set enrichment analysis

For GSEA analysis, we used gene set from Blood transcription modules from Li et al (35), bulk RNA sequencing of sorted tonsil B cells (Bm1+Bm2 and Bm5) and DZ and LZ GC gene lists from Victora et al. (36). We used clusterProfiler 3.8.1 to conduct GSEA analysis. Heatmaps displaying the signed -log10(FDR q value) of the result was created using GraphPad Prism (GraphPad Software, LLC).

### In vitro stimulation

Freshly isolated or thawed PBMC were washed with complete medium and counted. For qPCR, B cells were positively selected using anti-CD19 magnetic beads (Miltenyi Biotech). 0.5-1.0×10^6^ cells per well were seeded in round-bottom 96-well plates in 200μl 10%FBS/RPMI 1640. Cells were stimulated with anti-human IgM F(ab*′*2) or anti-human Ig(A+G+M) F(ab*′*2) (Jackson Immuno-Research Laboratories) for indicated times. After stimulation, cells were collected for flow cytometry or qPCR.

### Intracellular staining and Flow cytometry

To stain for flow cytometry, PBMC or tonsil cells were first incubated with Fc-Block (Biolegend) for 5 minutes at RT. Then 100ul of surface antibody cocktail (table S4) was added to each tube and incubated on ice for 30 minutes. Next, live/dead cell assay (Invitrogen) was applied for 15 minutes on ice. Intracellular staining was performed using Foxp3/Transcription factor staining buffer kit (eBioscience) following manufacture protocol. After intracellular staining, cells were washed and fixed in 100μl PBS/1% paraformaldehyde for 20 minutes at RT and then 200μl of PBS/5%BSA was added to dilute the fixative. Samples were analyzed on an 18-color LSR II flow cytometer (eBioscience) on the same day or after kept at 4°C overnight. Analyses were performed using FlowJo v.10 (Tree Star) and doublet exclusion was performed on all samples. Gating strategy is depicted in fig. S8.

### Immunofluorescent staining

List of primary and secondary antibodies used for immunofluorescent staining can be found in table S5. Detailed protocol is provided in supplementary materials.

### Statistical analysis

The statistical tests used are as described in each figure legend. P-values *≤* 0.05 are considered significant.

## Supporting information

all supplementary materials

## Supplementary Materials

### Materials and Methods

Fig. S1. Flow cytometry schema for sorting and analysis of B cells.

Fig. S2. Evidence of class-switched recombination and clonal expansion in RA synovium.

Fig. S3. NR4A+ cluster has characteristics consistent with tonsil intermediate zone B cells.

Fig. S4. Immunofluorescent staining of synovial tissue samples used for B cell single cell RNA sequencing.

Fig. S5. NR4A+ B cells in AMP RA cohort express chemokine receptors and cytokines important for ELS formation.

Fig. S6. A gene expression continuum from naïve to NR4A^+^ B cells.

Fig. S7. NR4A1, 2 and 3 mRNA expression are up-regulated in vitro through BCR stimulation.

Fig. S8. Flow cytometry schema for analysis of NR4A1+ B cells.

Table S1. Sample characteristics.

Table S2. List of overlapping genes between blood RA flare clusters and our single cell B cell clusters

Table S3. List of antibodies used for flow cytometry cell sorting

Table S4. List of antibodies used for flow cytometry

Table S5. List of antibodies used immunofluorescent staining

## Acknowledgements

We acknowledge the expertise and support of the University of Rochester Center for Musculoskeletal Research Histology Biochemistry and Molecular Imaging Core NIAMS AR069655 and the University of Rochester Resource Cores in Flow Cytometry and the Genomic Research Center.

## Author contributions

JHA, AM and NM conceived and designed the work. ED, DEO, SG and LTD were responsible for tissue sample acquisition and sample characterization. NM performed the sample processing, sorting, single-cell capture and flow cytometry experiments. JRM performed immunofluorescent staining of tissues. KER performed in vitro stimulation of B cells. JHA, AM and NM established the analytical strategies and analyzed the data. AM performed bioinformatic analyses. FZ, SR and the AMP network provided bioinformatic assistance. EC, EP, MB and CP provided tissue samples from the PEAC cohort. DEO provided analytic input and assistance with analysis of the RA flare cohort. JHA supervised the study. JHA, AM and NM wrote the manuscript. All authors contributed to reviewing and editing of the manuscript.

## Data and material availability

The raw data, gene expression matrices and cell annotations have been deposited in the Gene Expression Omnibus (https://www.ncbi.nlm.nih.gov/geo/) under accession number (pending). All other data needed to evaluate the conclusions in the paper are presented in the paper or the Supplementary Materials.

## Supplemental Acknowledgements

Accelerating Medicines Partnership Rheumatoid Arthritis & Systemic Lupus Erythematosus (AMP RA/SLE) Network member list

William Apruzzese^6^, Arnon Arazi^29,23^, Jennifer Barnard^1^, Jennifer Barnas^1^, Joan M. Bathon^14^, H. Michael Belmont^30^, Ami Ben-Artzi^15^, Celine Berthier^31^, Brendan F. Boyce^1^, David L. Boyle^16^, Michael B. Brenner^6^, S. Louis Bridges Jr^17^, Jill P. Buyon^30^, Vivian P. Bykerk^8^, Phillip Carlucci^30^, Arnold Ceponis^16^, Adam Chicoine^6^, Robert Clancy^30^, Sean Connery^32^, Carla M. Cuda^18^, Maria Dall’Era^33^, Anne Davidson^23^, Kevin Deane^19^, Wade DeJager^20^, Betty Diamond^23^, Kristina Deonaraine^30^, Salina Dominguez^18^, Patrick J. Dunn^21^, Thomas Eisenhaure^29^, Andrea Fava^34^, Andrew Filer^22^, Gary S. Firestein^16^, Lindsy Forbess^15^, Jennifer Goff^1^, Beatrice Goilav^35^, Ellen M. Gravallese^6^, Peter K. Gregersen^24^, Jennifer Grossman^37^, Joel M. Guthridge^21^, Nir Hacohen^29^, David Hildeman^37,38^, Jeffrey Hodgin^31^, V. Michael Holers^19^, Paul Hoover^29^, Diane Horowitz^23^, Raymond Hsu^33^, Laura B. Hughes^17^, Kazuyoshi Ishigaki^2,3,5^, Mariko Ishimori^15^, Peter Izmirly^30^, Judith A. James^20^, Tony Jones^29^, A. Helena Jonsson^6^, Kenneth C. Kalunian^16^, Diane L. Kamen^40^, Joshua Keegan^24^, Gregory Keras^6^, Ilya Korsunsky^2,3,4,5,6^, Matthias Kretzler^31^, Manjunath Kustagi^41^, Amit Lakhanpal^8^, James A. Lederer^6^, Myles Lewis^7^, Jessica Li^34^, Yuhong Li^6^, Zhihan Jian Li^6^, David Lieb^29^, Susan Macwana^20^, Holden Maecker^25,26^, Arthur M. Mandelin II^18^, Rong Mao^25,26^, Mandy J. McGeachy^27^, Maureen A. McMahon^36^, Joseph R. Mears^2,3,4,5^, Raji Menon^31^, Nghia Millard^2,3,4,5^, Larry Moreland^19^, Pavel Morozov^41^, Aparna Nathan^2,3,4,5^, Alessandra Nerviani^7^, Fernanda Payan-Schober^32^, Harris Perlman^18^, Michael Peters^29^, Michelle A. Petri^34^, Chaim Putterman^35^, Deepak A. Rao^6^, Karim Raza^22^, Christopher Ritchlin^1^, Felice Rivellese^7^, William H. Robinson^25,26^, Ilfita Sahbudin^22^, Saori Sakaue^2,3,4,5^, Karen Salomon-Escoto^28^, Jennifer Seifert^19^, Lorien Shakib^8^, Heather Sherman^8^, Daimon Simmons^6^, Anvita Singaraju^8^, Melanie Smith^8^, Hemant Suryawanshi^41^, Darren Tabechian^1^, Anjali Thakrar^18^, Thomas Tuschl^41^, Paul J. Utz^25,26^, Gerald Watts^6^, Kevin Wei^6^, Kathryn Weinand^2,3,4,5^, Michael Weisman^15^, David Wofsy^33^, E. Steve Woodle^39^, Qian Xiao^2,3,4,5^, Zhu Zhu^6^

^14^Division of Rheumatology, Department of Medicine, NewYork-Presbyterian/Columbia University Irving Medical Center, New York, NY, USA

^15^Division of Rheumatology, Cedars Sinai Medical Center, Los Angeles, CA, USA.

^16^Department of Medicine, Division of Rheumatology, Allergy and Immunology, University of California, San Diego, La Jolla, CA, USA.

^17^Division of Clinical Immunology and Rheumatology, Department of Medicine, University of Alabama at Birmingham, Birmingham, AL, USA.

^18^Division of Rheumatology, Department of Medicine, Northwestern University Feinberg School of Medicine, Chicago, IL, USA.

^19^Division of Rheumatology, University of Colorado School of Medicine, Aurora, CO, USA.

^20^Department of Arthritis & Clinical Immunology, Oklahoma Medical Research Foundation, Oklahoma City, OK, USA.

^21^ ImmPort Curation Team, NG Health Solutions, 2101 Gaither Road Rockville, MD 20850, USA.

^22^Rheumatology Research Group, Institute for Inflammation and Ageing, NIHR Birmingham Biomedical Research Center and Clinical Research Facility, University of Birmingham, Queen Elizabeth Hospital, Birmingham, UK.

^23^Institute of Molecular Medicine, Feinstein Institute for Medical Research, North Shore-LIJ Health System, Manhasset, NY, USA.

^24^Department of Surgery, Brigham and Women’s Hospital and Harvard Medical School, Boston, MA, USA.

^25^Division of Immunology and Rheumatology, Department of Medicine, Stanford University School of Medicine, Palo Alto, CA, USA.

^26^The Institute for Immunity, Transplantation, and Infection, Stanford University School of Medicine, Stanford, CA, USA.

^27^Division of Rheumatology and Clinical Immunology, University of Pittsburgh School of Medicine, Pittsburgh, PA, USA.

^28^Division of Rheumatology, Department of Medicine, University of Massachusetts Medical School, Worcester, MA, USA.

^29^Broad Institute of MIT and Harvard, Cambridge, MA, USA.

^30^Department of Medicine, Division of Rheumatology, New York University School of Medicine, New York, NY, USA.

^31^Internal Medicine, Department of Nephrology, University of Michigan, Ann Arbor, MI, USA.

^32^Department of Medicine, Paul L. Foster School of Medicine, Texas Tech University Health Sciences Center, El Paso, TX, USA.

^33^Rheumatology Division, University of California San Francisco, San Francisco, CA, USA.

^34^Division of Rheumatology, Johns Hopkins University, Baltimore, MD, USA.

^35^Division of Rheumatology and Department of Microbiology and Immunology, Albert Einstein College of Medicine and Montefiore Medical Center, Bronx, NY, USA.

^36^Department of Medicine, University of California Los Angeles, Los Angeles, CA, USA.

^37^Department of Pediatrics, University of Cincinnati, Cincinnati, OH, USA.

^38^Division of Immunobiology, Cincinnati Children’s Hospital Medical Center, Cincinnati, Ohio, USA.

^39^Division of Transplantation, Department of Surgery, University of Cincinnati College of Medicine, Cincinnati, OH, USA.

^40^Division of Rheumatology and Immunology, Medical University of South Carolina, Charleston, SC, USA.

^41^Laboratory of RNA Molecular Biology, Rockefeller University, New York, NY, USA.

