## Supplementary material for "Single cell analysis of RA synovial B cells reveals a dynamic spectrum of ectopic lymphoid B cell activation and hypermutation characterized by NR4A nuclear receptor expression": all supplementary materials

Supplemental material

#### Materials and Methods

##### Quantitative real-time polymerase chain reaction (qPCR)

B cells were collected in RLTplus lysis buffer (Qiagen) supplemented with 1% 2-mercaptoethanol, and RNA were extracted using *Quick*-RNA kit (Zymo Research) and concentration was measured by NanoDrop (Thermo Scientific). Complementary DNA (cDNA) synthesis was done using a qScript cDNA synthesis kit (Quantabio). TaqMan expression assay system for NR4A1 (Hs00374226_m1), NR4A2 (Hs01117527_g1), NR4A3 (Hs00545009_g1) from Applied Biosystems were used to detect NR4A1, NR4A2 and NR4A3 mRNA expression, respectively. PPIA (Hs04194521_s1, Applied Biosystems) was used as a control housekeeping gene. The fold change in mRNA expression relative to control was calculated using the comparative threshold cycle ($\Delta\Delta C_t$) method.

##### Immunofluorescent staining

Primary antibodies: List of primary and secondary antibodies used for immunofluorescent staining can be found in table S5. 5 μm formalin-fixed paraffin sections were incubated at 60ºC overnight (ON). Tissue sections were quickly transferred to xylenes and gradually hydrated by transferring slides to absolute alcohol, 96% alcohol, 70% alcohol, and water. Slides were immersed in an antigen retrieval solution, boiled for 30 minutes, and cooled down for 10 minutes at room temperature (RT). Slides were rinsed several times in water and transferred to PBS. Non-specific binding was blocked with 5% normal donkey serum in PBS containing 0.1% Tween 20, 0.1% Triton-X-100 for 30 minutes, at RT in a humid chamber. Primary antibodies were added to slides and incubated in a humid chamber at room RT, ON. Slides were quickly washed in PBS, and fluorescently labeled, secondary antibodies were incubated for 2 hours at RT in a humid chamber. Finally, slides were rinsed for 1 hour in PBS and mounted with Vectashield antifade mounting media with DAPI (H-1200, Vector Laboratories). 200x and 200x 3x3 mosaic pictures were taken with a Zeiss Axioplan 2 microscope and recorded with a Hamamatsu camera.

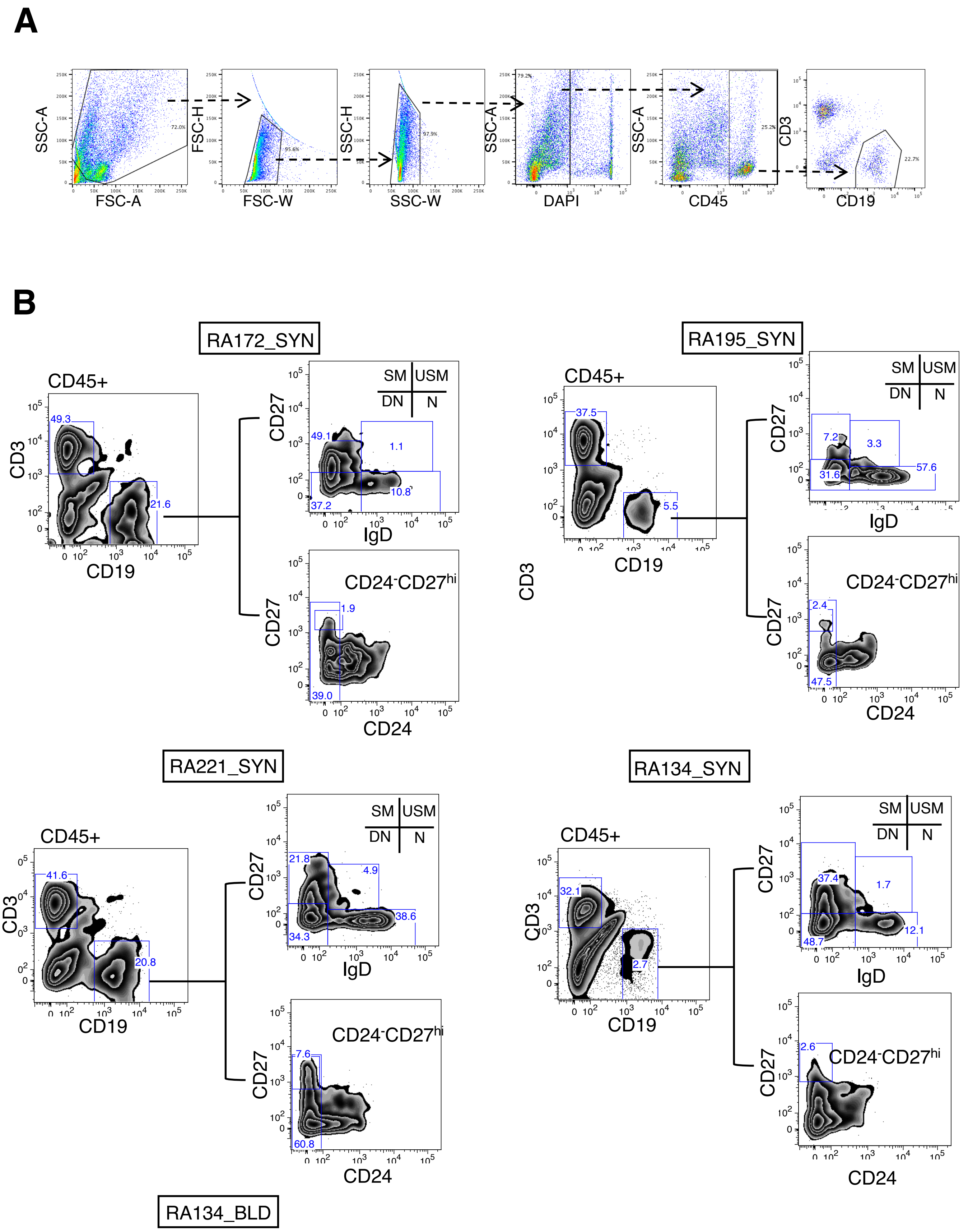


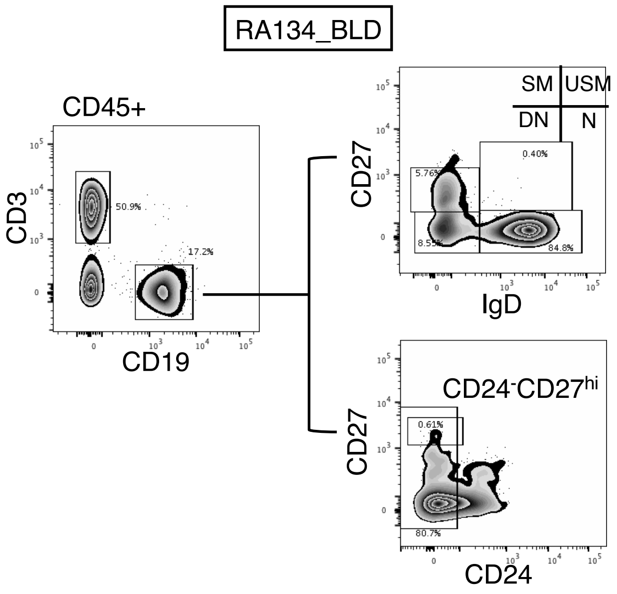


**fig. S1**. Flow cytometry schema for sorting and analysis of B cells. **(A)** Flow cytometry gating strategy for sorting of B cells. **(B)** Flow cytometry analysis for B cell phenotyping for each sample. CD27 vs. IgD plot shows % of switched memory (SM), un-switched memory (USM), double negative (DN) and Naïve (N). CD27 vs. CD24 identify $CD24^{-}CD27^{hi}$ plasmablast.

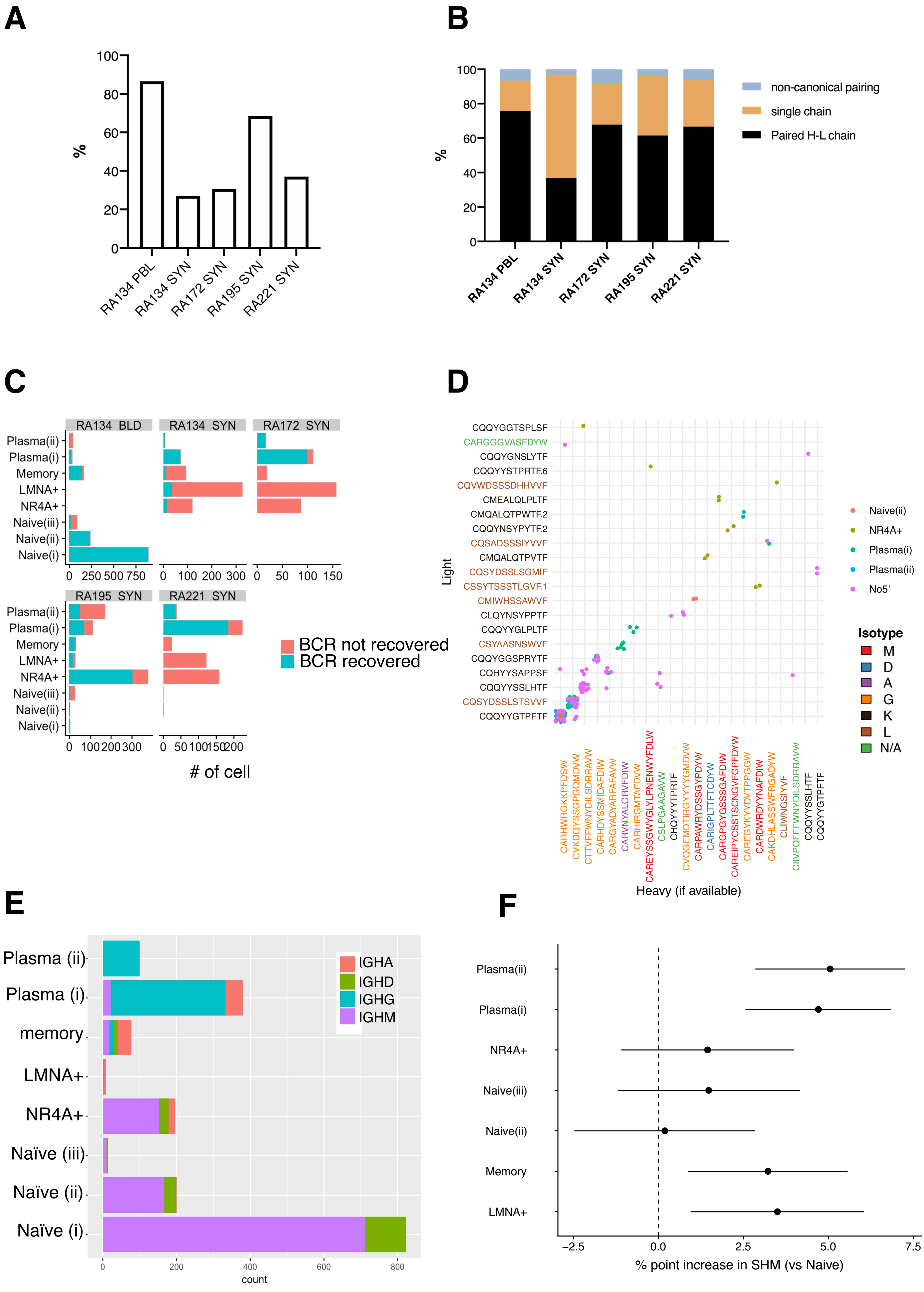

**fig. S2**. **Evidence of class-switched recombination and clonal expansion in RA synovium. (A)** Bar graph showing percentage of cells with productive V-J spanning pair out of total barcode recovered per sample, RA134 PBL (1559 cells), RA134 SYN (694 cells), RA172 SYN (457 cells), RA195 SYN (1240 cells) and RA221 SYN (607 cells). **(B)** Bar graph displaying percentage recovery of paired heavy-light chain, single heavy or light chain or non-canonical pairing out of total, RA134 PBL (1348 cells), RA134 SYN (187 cells), RA172 SYN (140 cells), RA195 SYN (850 cells) and RA221 SYN (225 cells). **(C)** Bar graph showing number of cells where BCR was recovered or not recovered per cluster and per sample. **(D)** Pairing of heavy and light chains in RA195 SYN. Each point represents a cell. Clones that occurred 2 or more times are shown. The amino acid sequence of the CDR3 of the heavy and light chains are shown below and the left of the x and y axes, respectively. The color of the label represents the isotype. **(E)** Bar graphs depicting number of cells expressing IGHA, IGHD, IGHG and IGHM per B cell cluster. **(F)** Difference in SHM rate in cell populations vs Naive(i) using linear mixed effect models with capture:cluster random effect.

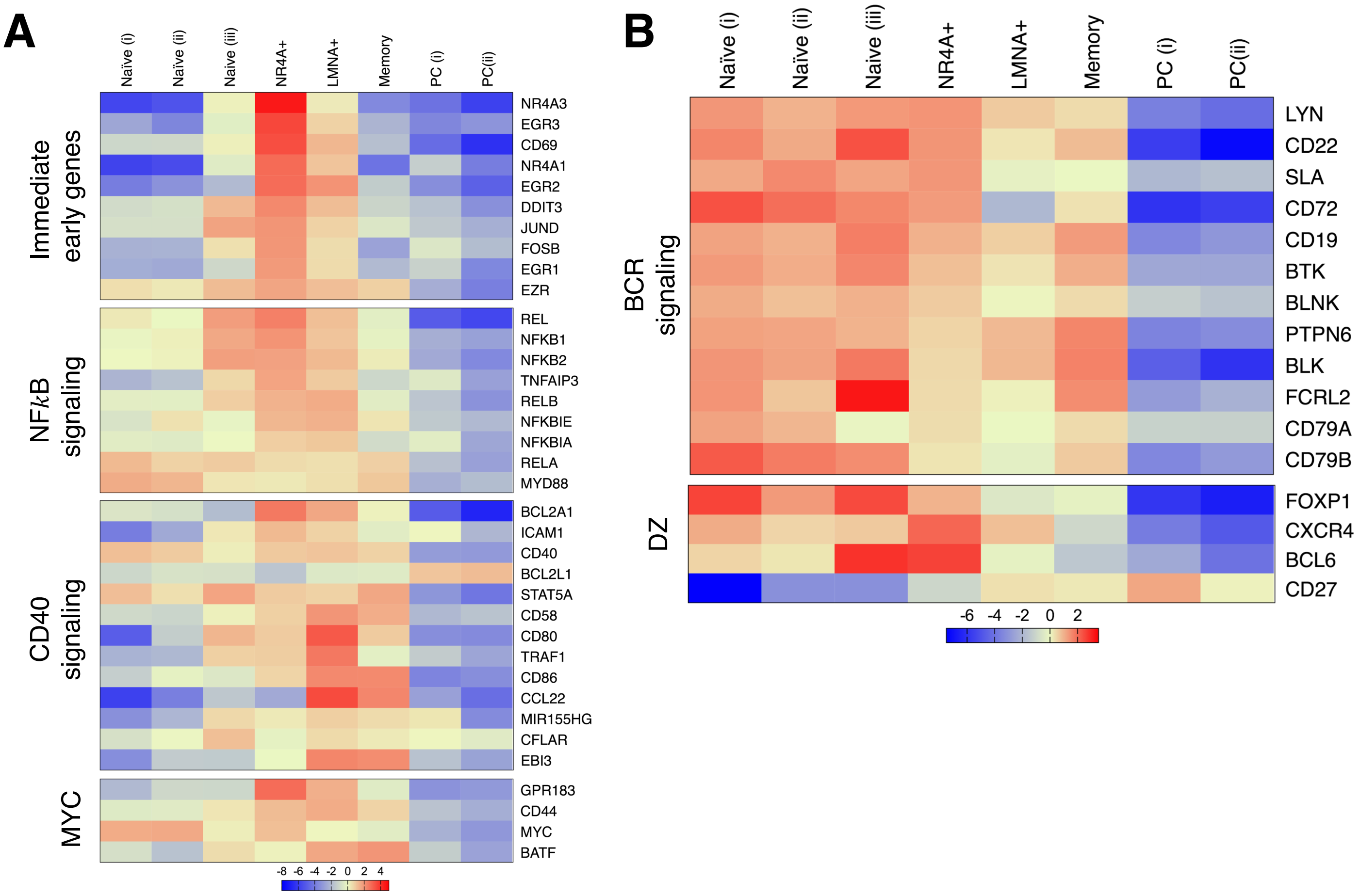


**fig. S3. NR4A+ cluster has characteristics consistent with tonsil intermediate zone B cells. (A)** Heatmap display log2 fold change of immediate early genes, NF*k*B signaling genes, CD40 signaling genes and MYC genes (Ye et al, 2010; Ollila et al, 2003; Holmes et al, 2020). **(B)** NR4A⁺ cluster maintains expression of some DZ genes. Heatmap display log2 fold change of BCR signaling genes and genes associated with DZ GC B cells in each B cell cluster (Holmes et al, 2020).

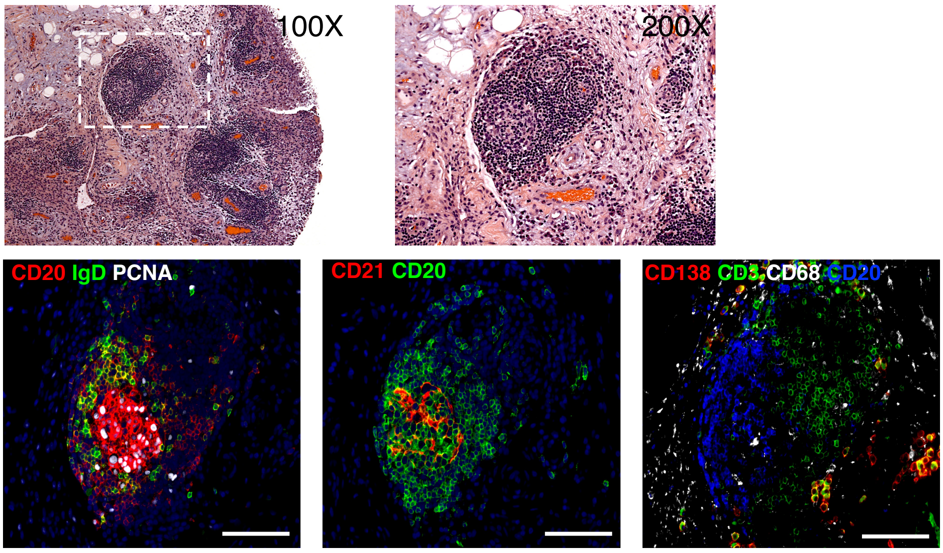


**fig. S4**. Immunofluorescent staining of synovial tissue samples used for B cell single cell RNA sequencing. 5 μm synovial serial sections were stained with Hematoxylin and Eosin or antibodies specific for proliferating cell nuclear antigen (PCNA, white), IgD (green), and CD20 (red) to detect large $CD20^{+}IgD^{-}PCNA^{+}$ germinal center B cells. Synovial sections were stained with a combination of antibodies specific for CD20 and CD21 to detect $CD21^{+}$ follicular dendritic cells (red) in germinal centers populated by $CD20^{+}$ B cells (green). We also stain paraffin sections with antibodies to detect $CD138^{+}$ plasma cells (red), $CD3^{+}$ T cells (green), $CD68^{+}$ macrophages (white), and $CD20^{+}$ B cells (blue) in the ELS in the inflamed synovia of RA patient. Representative 200x magnification pictures are shown. Scale bars represent 100 μm.

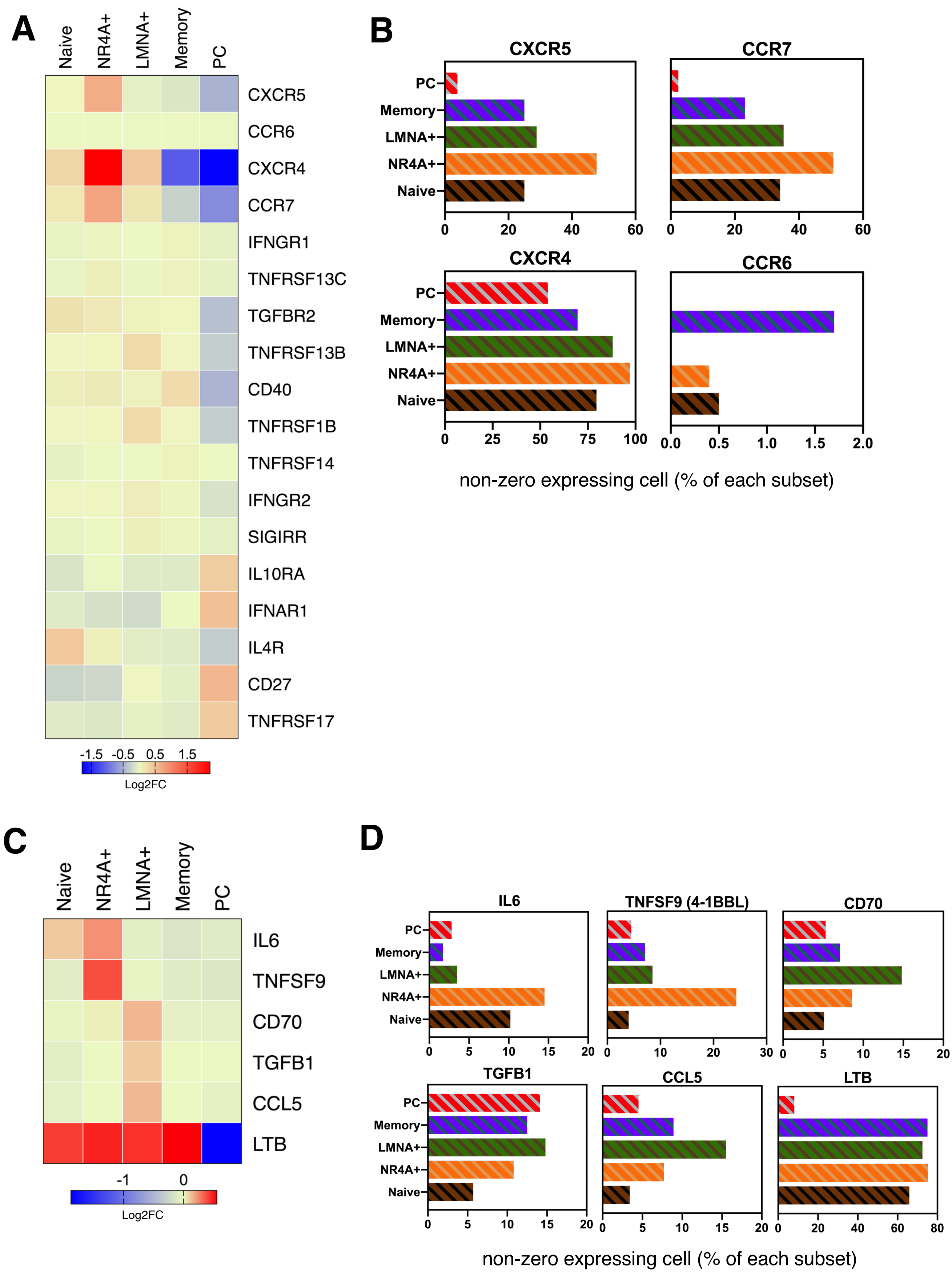


**fig. S5. NR4A+ B cells in AMP RA cohort express chemokine receptors and cytokines important for ELS formation.** Single cell RNA sequencing data of synovial B cells from AMP phase I (n= 14 RA, Zhang et al., 2019) were re-analyzed and B cell subsets parallel to current study were identified using a supervised classification technique SingleR. **(A)** Heatmap displays log2 fold change of cytokine and chemokine receptors in each B cell subset. **(B)** Bar graphs display percentage of cells of each subset that express *CXCR5, CCR7, CXCR4* or *CCR6*. **(C)** Heatmap displays log2 fold change of cytokines and chemokines in each B cell subset. **(D)** Bar graph represent percentage of cells of each subset that express *IL6, TNFSF9, CD70, TGFB1, CCL5* or *LTB*.

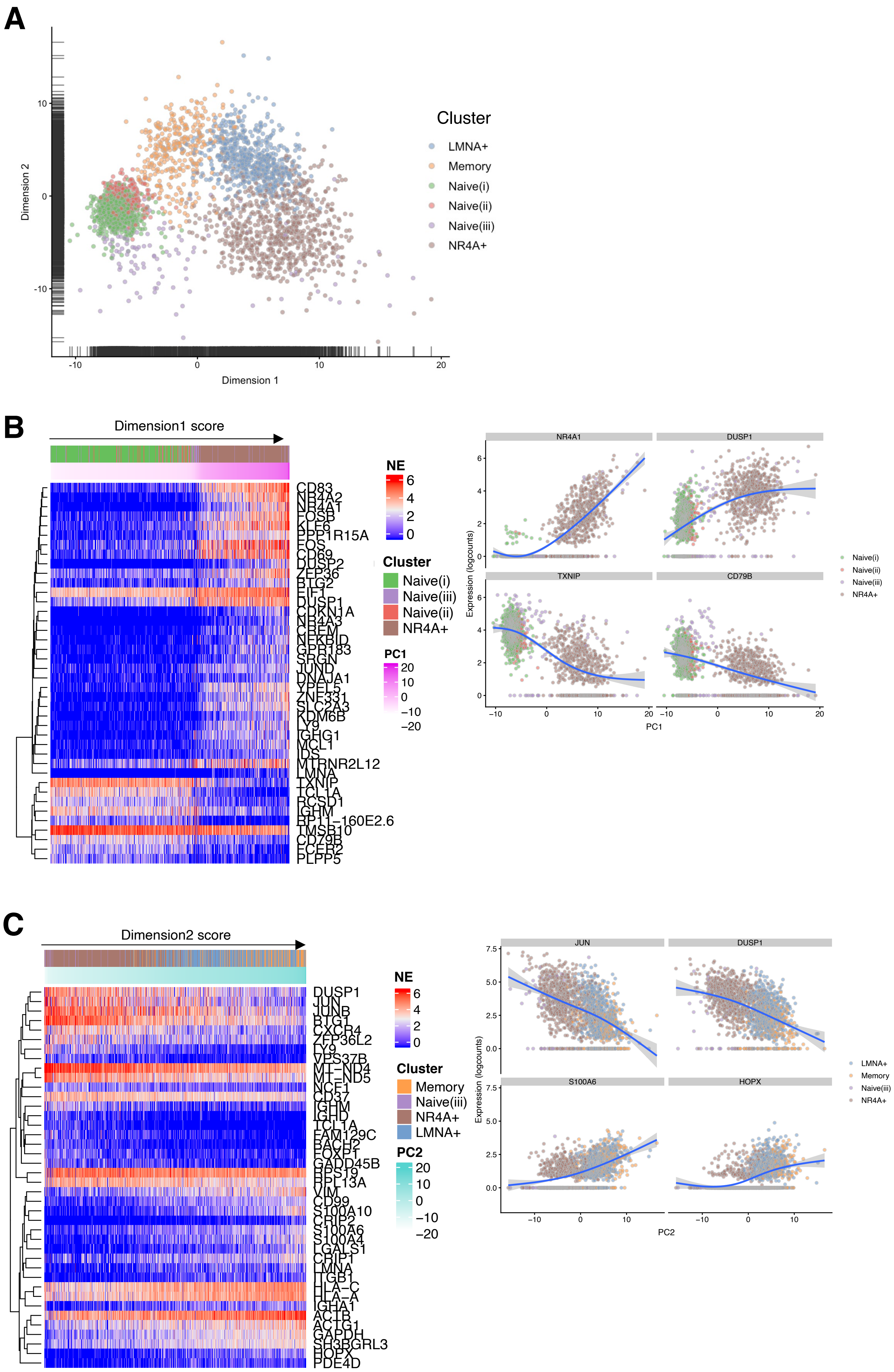
A
Dimension 1
Dimension 2
Cluster
LMNA+
Memory
Naive(i)
Naive(ii)
Naive(iii)
NR4A+
B
Dimension1 score
NE
Cluster
Naive(i)
Naive(iii)
Naive(ii)
NR4A+
PC1
CD83
NR4A2
NR4A1
FOSB
KLF6
PPP1R15A
FOS
CD69
DUSP2
ZFP36
BTG2
EIF1
DUSP1
CDKN1A
NR4A3
CREM
NFKBID
GPR183
SRGN
JUND
DNAJA1
YPEL5
ZNF331
SLC2A3
KDM6B
LY9
IGHG1
MCL1
IDS
MTRNR2L12
LMNA
TXNIP
TCL1A
RCSD1
IGHM
RP11-160E2.6
TMSB10
CD79B
FCER2
PLPP5
NR4A1
DUSP1
TXNIP
CD79B
Expression (logcounts)
PC1
C
Dimension2 score
NE
Cluster
Memory
Naive(iii)
NR4A+
LMNA+
PC2
DUSP1
JUN
JUNB
BTG1
CXCR4
ZFP36L2
LY9
VPS37B
MT-ND4
MT-ND5
NCF1
CD37
IGHM
IGHD
TCL1A
FAM129C
BACH2
FOXP1
GADD45B
RPS19
RPL13A
VIM
CD99
S100A10
CRIP2
S100A6
S100A4
LGALS1
CRIP1
LMNA
ITGB1
HLA-C
HLA-A
IGHA1
ACTB
ACTG1
GAPDH
SH3BGRL3
HOPX
PDE4D
JUN
DUSP1
S100A6
HOPX
Expression (logcounts)
PC2

**D**

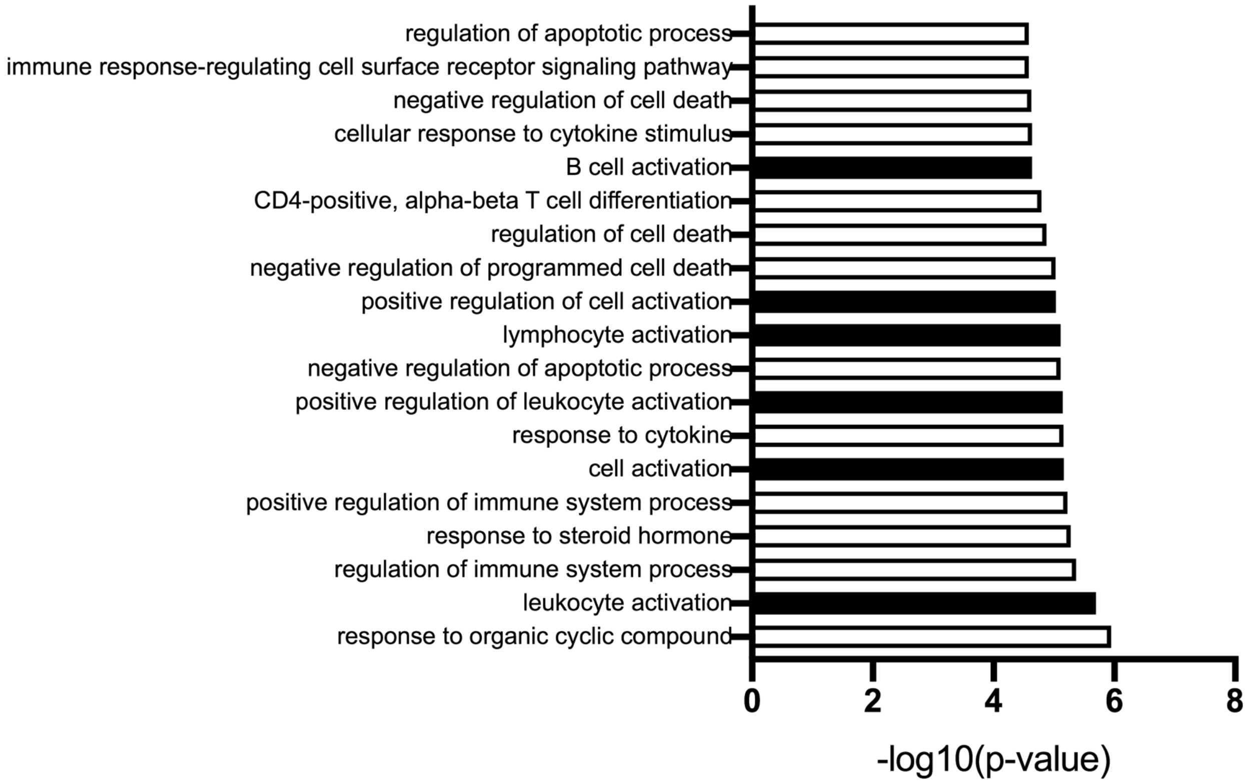


**fig. S6. A gene expression continuum from naïve to NR4A⁺ B cells. (A)** Principal component analysis projecting gene expression data of Naïve (i), Naïve (ii), Naïve (iii), NR4A⁺, LMNA⁺ and memory B cell subsets onto two dimensional space. **(B)** Heatmap display expression of top 40 loading genes for dimension 1 with increasing score from Naïve (i), Naïve (ii), Naïve (iii) and NR4A⁺ subset. Plots on the right show example of genes increase (NR4A1, DUSP1) or decrease (TXNIP, CD79B) as cells transition from naïve to NR4A⁺ state. **(C)** Heatmap display expression of top 40 loading genes for dimension 2 with increasing score from Naïve (iii), NR4A⁺, LMNA⁺ and memory subset. Plots on the right show example of genes decrease (JUN, DUSP1) or increase (S100A6, HOPX) as cells transition from NR4A⁺ to memory state. Color on the heatmap from blue to red represent normalized expression from low to high. The gradient of pink color shows gene loading score along dimension 1 (PC1) or dimension 2 (PC2). **(D)** Gene list enrichment analysis was done using ToppGene Suite by inputting 486 differential expressed genes of NR4A+ cluster. Top 19 of significant GO: Biological processes are shown. Black bars indicate biological processes relate to cell activation.

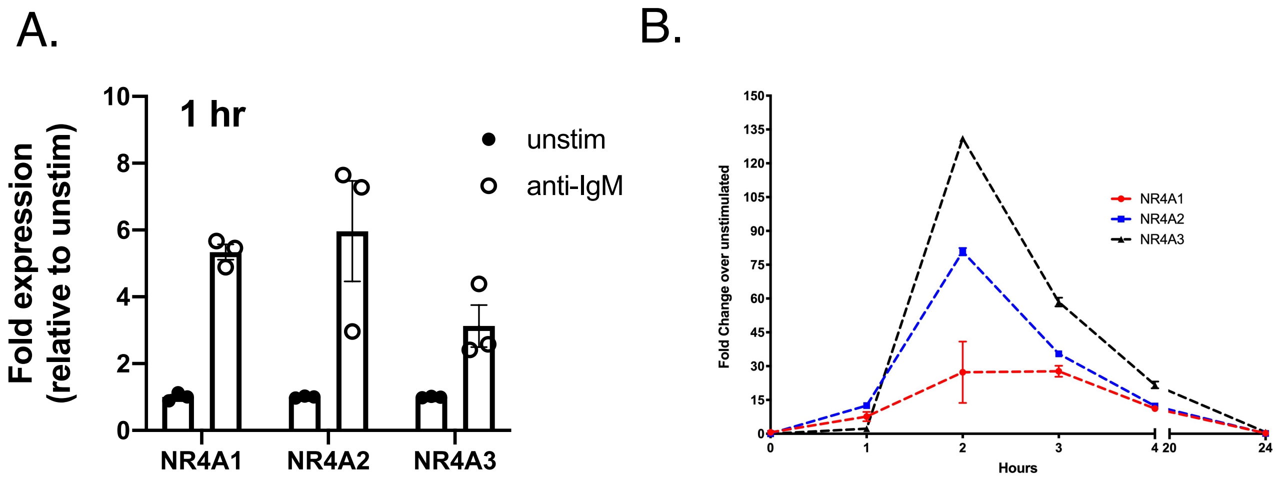


**fig. S7**. **NR4A1, 2 and 3 mRNA expression is up-regulated in vitro through BCR stimulation. (A**) B cells were purified from PBMC of healthy donors (n=3) cultured with or without anti-IgM (20$\mu$M) for 1 hour. Quantitative real-time PCR was performed for NR4A1, NR4A2, NR4A3 and housekeeping gene control PPIA. Fold expressions were calculated using ΔΔCt method relative to unstimulated control. (**B**). B cells were stimulated with anti-IgM for 1, 2 , 3, 4 and 24 hours. Fold expression changes for NR4A1, 2 and 3 were calculated relative to unstimulated control.

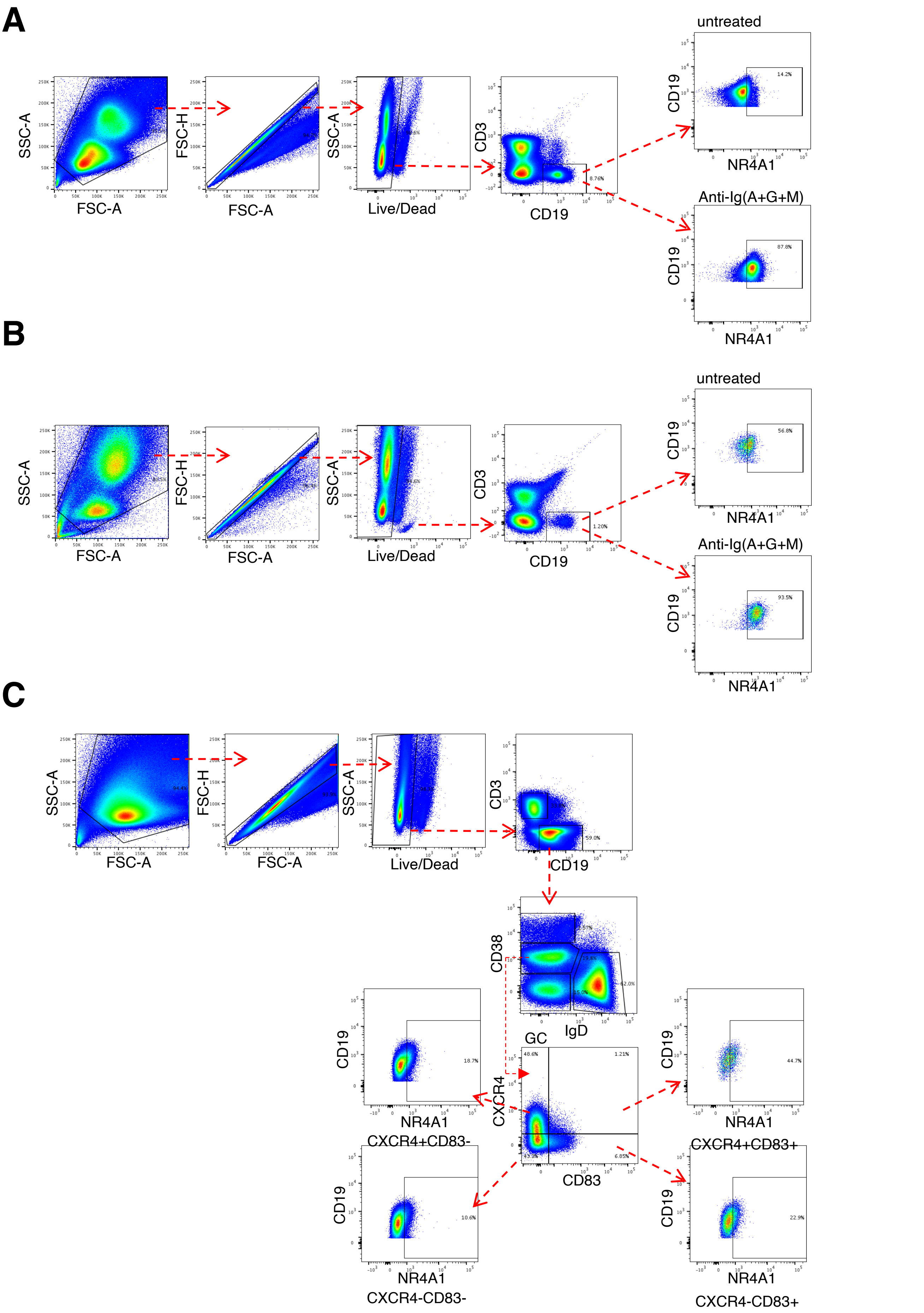


**fig. S8. Flow cytometry schema for analysis of NR4A1+ B cells. (A)** Flow cytometry gating strategy for PBMC. Doublet exclusion was performed, followed by dead cell exclusion. NR4A1+ cells were gated within CD19+ B cell. Representative dot plots from untreated and anti-Ig(A+G+M) treated from the same PBMC sample are shown.**(B)** Flow cytometry gating strategy for synovial fluid. Doublet exclusion was performed, followed by dead cell exclusion. NR4A1+ cells were gated within CD19+ B cell. Representative dot plots from untreated and anti-Ig(A+G+M) treated from the same synovial fluid sample are shown. **(C)** Gating strategy for tonsil B cells. After doublet and dead cell exclusion, CD19+ B cells were further characterized based on CD38 vs. IgD expression and germinal center B cells (GC) were defined as IgD-CD38+. GC B cells were further subset using CXCR4 and CD83 expression and NR4A1+ cells were determined within each subset.

**Table S1**: Sample characteristics.

| **RA I.D.** | **RA172** | **RA195** | **RA221** | **RA134** |
|---|---|---|---|---|
| Anti-CCP | negative | positive | nd | positive |
| DAS28 | 4.79 | 4.79 | nd | 5.8 |
| Inflammation score* | 3 | 3 | 4 | 3 |
| Plasma cell score** | 2 | 1 | 2 | 1 |
| Lining score*** | 2 | 2 | 3 | 2 |

The inflammation, plasma cells and lining scores were evaluated by a board certified pathologist.

*Synovial Lymphocytic Inflammation: Mag. 4x or 5x: (0) –None, (1) – Mild (0-1 perivascular aggregates per low power field), (2) – Moderate (>1 perivascular aggregate + focal interstitial infiltration), (3) – Marked (both perivascular and widespread interstitial aggregates), (4) – Band-like (extensive lymphocyte presence across tissue)

** Plasma Cell Inflammation (x25mag): (0) – <10% plasma cells within lymphocytic aggregates, (1) – <50% plasma cells, (2) – >50% plasma cells.

***Synovial Lining Hyperplasia: Mag. 10x or 20x: (0) – Normal Lining, (1) – 1-3 Cells Thick, (2) – 3-4 Cells Thick, (3) – >4 Cells Thick.

nd = not determined

### **Table S2**: List of overlapping genes between blood RA flare clusters and our single cell B cell clusters

| Cluster | syno_seurat_cluster | prime_cluster | Description | GeneRatio | BgRatio | pvalue | p.adjust | qvalue | geneID | Count | log10_fdr |
|---|---|---|---|---|---|---|---|---|---|---|---|
| LMNA+ | LMNA+ | 1 | 1 | 19/286 | 598/29420 | 7.38E-06 | 3.69E-05 | 7.76E-06 | KLF6/FTH1/VIM/CD44/MCL1/YWHAZ/CD53/LRRFIP1/LITAF/PELI1/ATP6V0E1/GNG2/WIPF1/CAST/FAM49A/RAB31/TMX4/CDC42EP3/AIM2 | 19 | 5.10994618 |
| LMNA+ | LMNA+ | 2 | 2 | 16/286 | 542/29420 | 9.24E-05 | 2.31E-04 | 4.86E-05 | CD74/RPLP2/RPS15/PFN1/SH3BGRL3/LMNA/PPDPF/PLP2/ZYX/EMP3/HIGD2A/MYADM/C9orf16/FAM102A/ZBTB32/C4orf48 | 16 | 4.31325515 |
| LMNA+ | LMNA+ | 5 | 5 | 5/286 | 151/29420 | 0.01627393 | 0.02712321 | 0.00571015 | RPS29/CD83/RPL38/ODC1/UBB | 5 | 2.24335252 |
| LMNA+ | LMNA+ | 4 | 4 | 6/286 | 249/29420 | 0.0354861 | 0.04435762 | 0.00933845 | RPLP1/RPL36/BCAS4/NR4A1/CRIP2/CHCHD10 | 6 | 2.02972538 |
| LMNA+ | LMNA+ | 3 | 3 | 2/286 | 408/29420 | 0.90854387 | 0.90854387 | 0.19127239 | FOSB/EGR1 | 2 | 0.71834771 |
| Memory | Memory | 2 | 2 | 19/224 | 542/29420 | 4.16E-08 | 1.67E-07 | 8.76E-08 | CD74/RPLP2/SH3BGRL3/HLA-DRB1/PFN1/MT-CO1/PTPRCAP/EMP3/PLP2/PPDPF/GPSM3/ZYX/C9orf16/PGLS/C4orf48/ZBTB32/AES/SIGIRR/HLA-G | 19 | 7.05733241 |
| Memory | Memory | 1 | 1 | 10/224 | 598/29420 | 0.0168605 | 0.03372101 | 0.0177479 | CD53/VIM/RHOA/AIM2/LCP1/LITAF/ATP6V0E1/TMEM154/RAB31/CAST | 10 | 1.75085303 |
| Memory | Memory | 4 | 4 | 4/224 | 249/29420 | 0.12306012 | 0.16408015 | 0.08635798 | RPL36/RPLP1/BLK/CRIP2 | 4 | 1.06369755 |
| Memory | Memory | 5 | 5 | 1/224 | 151/29420 | 0.6855898 | 0.6855898 | 0.36083674 | RPS29 | 1 | 0.44268925 |
| Naive(i) | Naive(i) | 2 | 2 | 33/456 | 542/29420 | 3.16E-11 | 1.26E-10 | 3.33E-11 | CD37/HLA-DRB1/CD74/IER2/CD79A/RPL18/RPS15/IGHD/RPL3/PCBP1/EEF2/AES/YBX3/KLF2/IFITM3/PTPRCAP/GPSM3/LBH/PLD4/C16orf74/PAX5/H2AFX/MAF1/TAPBP/SPI1/DBNDD1/VPS51/BANF1/JUP/H1FX/HLA-DOA/SIPA1L3/DRAP1 | 33 | 10.4781199 |
| Naive(i) | Naive(i) | 1 | 1 | 21/456 | 598/29420 | 4.87E-04 | 9.73E-04 | 2.56E-04 | SELL/ITM2B/LTA4H/MYLIP/KIAA0040/RHOA/LBR/YWHAB/LYN/EIF4G2/SNX10/FLI1/SSH2/SP110/LYST/AFF1/PRCP/SRPK2/TBC1D1/IFNGR2/CYBB | 21 | 3.59156483 |
| Naive(i) | Naive(i) | 4 | 4 | 10/456 | 249/29420 | 0.00576935 | 0.00769247 | 0.00202433 | TCL1A/RPL18A/FCER2/FAM129C/RPL36/BCL7A/SNX22/BLK/LIMS2/TLE1 | 10 | 2.69371783 |
| Naive(i) | Naive(i) | 5 | 5 | 2/456 | 151/29420 | 0.6814974 | 0.6814974 | 0.17934142 | RPS19/SNHG7 | 2 | 0.74631939 |
| Naive(ii) | Naive(ii) | 2 | 2 | 20/241 | 542/29420 | 2.65E-08 | 1.06E-07 | 8.37E-08 | CD74/HLA-DRB1/RPL18/RPS15/CD37/CD79A/RPL3/EEF2/PTPRCAP/IER2/IGHD/PCBP1/IFITM3/PLD4/C16orf74/AES/GPSM3/RNF187/DRAP1/APRT | 20 | 7.0773004 |
| Naive(ii) | Naive(ii) | 4 | 4 | 4/241 | 249/29420 | 0.1486277 | 0.29447137 | 0.23247739 | RPL18A/TCL1A/FCER2/FAM129C | 4 | 0.63361927 |
| Naive(ii) | Naive(ii) | 1 | 1 | 7/241 | 598/29420 | 0.22085352 | 0.29447137 | 0.23247739 | ITM2B/SELL/LTA4H/KIAA0040/PGK1/IFNGR2/ARPC5 | 7 | 0.63361927 |
| Naive(ii) | Naive(ii) | 5 | 5 | 2/241 | 151/29420 | 0.35124456 | 0.35124456 | 0.27729834 | RPS19/SNHG7 | 2 | 0.55705273 |
| Naive(iii) | Naive(iii) | 1 | 1 | 87/1539 | 598/29420 | 4.77E-18 | 2.38E-17 | 1.51E-17 | TMEM2/MDM4/DIP2B/CELF2/TRIM38/RC3H1/RBM5/SIPA1L1/SON/FCHSD2/MCTP2/LYST/KMT2C/UBR2/IQGAP1/CLMN/SSH2/MAP3K8/AVL9/NSF/TACC1/NUMB/SPTLC2/KAT6A/NECAP1/ARHGAP24/MYLIP/ELOVL5/U2AF1/TBC1D1/NPEPPS/CDK14/SLC12A6/LRRK2/CLTC/TMED8/TET2/CCDC6/AP1G1/EIF2AK2/RNF145/SBNO | 87 | 16.8221174 |
| Naive(iii) | Naive(iii) | 2 | 2 | 40/1539 | 542/29420 | 0.01862043 | 0.04655106 | 0.02940067 | MT-CO1/IGHM/IGHD/YBX3/SIPA1L3/RBM38/PAX5/TCF3/HCFC1/KDM4B/PITPNM1/CTBP1/NUMA1/TTC7A/MXD4/UBE2O/JUND/MKNK2/CLEC16A/PER1/PPP1R9B/IER5/USF2/AP2A1/ASCC2/PPP6R1/HPS1/CABIN1/SPSB3/HLA-G/ALDH16A1/AES/RNF44/ZSCAN18/TGFB1/SCAMP3/DCXR/TSC22D4/KLF2/BAG6 | 40 | 1.53164276 |
| Naive(iii) | Naive(iii) | 4 | 4 | 12/1539 | 249/29420 | 0.65581481 | 0.99999999 | 0.63157894 | FAM129C/QSOX2/TCL1A/BLK/BCL7A/TLE1/CCS/WHAMM/FCER2/TNFRSF13C/QTRT1/SNX22 | 12 | 0.19957236 |
| Naive(iii) | Naive(iii) | 5 | 5 | 5/1539 | 151/29420 | 0.90123068 | 0.99999999 | 0.63157894 | ABCB4/OSBPL10/RASGRP3/AFG3L2/CD83 | 5 | 0.19957236 |
| Naive(iii) | Naive(iii) | 3 | 3 | 2/1539 | 408/29420 | 0.99999999 | 0.99999999 | 0.63157894 | MATN1/FOSB | 2 | 0.19957236 |
| NR4A+ | NR4A+ | 1 | 1 | 24/342 | 598/29420 | 1.77E-07 | 8.84E-07 | 3.72E-07 | SLC2A3/KLF6/FTH1/MCL1/MAP3K8/YWHAZ/CD44/GNG2/NFKBIZ/NFKB1/SDCBP/PPP1R15B/LRRFIP1/LYN/STK4/TMEM2/ARHGAP24/PRKAR1A/WIPF1/NR3C1/BCL6/GNA13/LITAF/HSPA1A | 24 | 6.42912544 |
| NR4A+ | NR4A+ | 2 | 2 | 15/342 | 542/29420 | 0.00186185 | 0.00465463 | 0.00195984 | JUNB/JUND/RPLP2/MT-CO1/CDKN1A/MYADM/PER1/KLF2/MARCKSL1/KLF4/MIDN/TUBB2A/NRROS/BHLHE40/LMNA | 15 | 2.7077789 |
| NR4A+ | NR4A+ | 5 | 5 | 6/342 | 151/29420 | 0.00861865 | 0.01436441 | 0.00604817 | CD83/RPS29/RPL38/SNHG8/ODC1/HIST1H4C | 6 | 2.21837571 |
| NR4A+ | NR4A+ | 4 | 4 | 5/342 | 249/29420 | 0.16551997 | 0.20689996 | 0.08711577 | NR4A1/CXCR5/HNRNPA0/STAG3/ID3 | 5 | 1.05990321 |
| NR4A+ | NR4A+ | 3 | 3 | 2/342 | 408/29420 | 0.95191929 | 0.95191929 | 0.40080812 | FOSB/EGR1 | 2 | 0.39706349 |
| Plasma(i) | Plasma(i) | 4 | 4 | 8/231 | 249/29420 | 8.30E-04 | 0.00332167 | 0.00174825 | MZB1/IGLC2/IGKV1-5/IGLL5/IGKV1-6/IGKV4-1/IGLV3-19/IGHV3-11 | 8 | 2.75739711 |
| Plasma(i) | Plasma(i) | 1 | 1 | 11/231 | 598/29420 | 0.00800738 | 0.01601475 | 0.00842882 | PRDM1/FNDC3B/SEL1L/NEAT1/DNAJC3/IFNAR2/UBE2J1/FUCA2/CCPG1/VIM/HSPA1A | 11 | 2.07423341 |
| Plasma(i) | Plasma(i) | 2 | 2 | 8/231 | 542/29420 | 0.06525558 | 0.08700744 | 0.04579339 | DERL3/ITM2C/ST6GALNAC4/PRDX2/CST3/IGHM/CDK2AP2/IGHA1 | 8 | 1.33919719 |
| Plasma(i) | Plasma(i) | 3 | 3 | 1/231 | 408/29420 | 0.96078601 | 0.96078601 | 0.50567685 | DNAAF1 | 1 | 0.29612693 |
| Plasma(ii) | Plasma(ii) | 4 | 4 | 9/126 | 249/29420 | 1.34E-06 | 4.03E-06 | 2.83E-06 | MZB1/IGKV4-1/IGLC2/IGLV3-19/IGLL5/IGKV1-5/IGHV3-11/TECR/IGKV1-6 | 9 | 5.54858543 |
| Plasma(ii) | Plasma(ii) | 2 | 2 | 4/126 | 542/29420 | 0.20338045 | 0.30507067 | 0.21408468 | DERL3/ITM2C/CST3/IGHA1 | 4 | 0.66941441 |
| Plasma(ii) | Plasma(ii) | 1 | 1 | 2/126 | 598/29420 | 0.72883856 | 0.72883856 | 0.51146565 | FNDC3B/SEL1L | 2 | 0.29118352 |

**Table S3**: List of antibodies used for flow cytometry cell sorting

| Antigen | Fluorochrome | Clone | Vendor | Catalog # |
|---|---|---|---|---|
| Pdn | PE | NZ-1.3 | eBiosciences | 12-9381-42 |
| CD90 | PerCP-cy5.5 | 5E10 | BD Biosciences | 561557 |
| CD45 | Brilliant Violet 786 | HI30 | BD Biosciences | 563718 |
| CD14 | Alexa Fluor 700 | M5E2 | BD Biosciences | 557923 |
| IgD | FITC | IA6-2 | BD Biosciences | 555778 |
| CD19 | PE/Dazzle 594 | HIB19 | Biolegend | 302251 |
| CD3 | PE-cy5 | UCHT1 | BD Biosciences | 555334 |
| CD27 | Brilliant Violet 605 | O323 | Biolegend | 302830 |
| CD4 | APC-cy7 | OKT4 | Biolegend | 317418 |
| CD11c | PE-cy7 | 3.9 | Biolegend | 301608 |
| CD24 | Alexa Fluor 647 | ML5 | Biolegend | 311109 |

**Table S4**: List of antibodies used for flow cytometry

| Antigen | Fluorochrome | Clone | Vendor | Catalog # |
|---|---|---|---|---|
| CD19 | APC-cy7 | SJ25C1 | BD Biosciences | 557791 |
| CD3 | Alexa Fluor 700 | UCH1 | Biolegend | 300424 |
| CD27 | Brilliant Violet 605 | O323 | Biolegend | 302830 |
| IgD | BUV737 | IA6-2 | BD Biosciences | 612798 |
| CD38 | PerCP-cy5.5 | HIT2 | BD Biosciences | 551400 |
| NR4A1 | PE | 12.14 | ThermoFisher Scientific | 12-5965-82 |
| GPR183 | Brilliant Violet 421 | 1C6 | BD Biosciences | 562558 |
| BCL6 | PE/Dazzle 594 | 7D1 | Biolegend | 358510 |
| CD69 | PE-cy5 | FN50 | BD Biosciences | 555532 |
| CD83 | Alexa Fluor 647 | HB15e | Biolegend | 305314 |
| IgG | Brilliant Violet 786 | G18-145 | BD Biosciences | 564230 |
| Ki67 | BUV395 | B56 | BD Biosciences | 564071 |
| CD77 | FITC | 5B5 | BD Biosciences | 551353 |
| CXCR4 | PE-cy7 | 12G5 | BD Biosciences | 560669 |

**Table S5**: List of antibodies used immunofluorescent staining

| Antibody | Clone/Fluorochrome | Vendor | Catalog # |
|---|---|---|---|
| **Primary AB** | | | |
| goat anti-human CD20 | polyclonal | LifeSpan Biosciences | LS-B11144 |
| rabbit anti- NR4A1 | polyclonal | Sigmaaldrich | HPA070142 |
| mouse anti-human CD21 | 2G9 | Thermo Fisher Scientific | MA5-11417 |
| mouse anti-human Ki67 | MIB-1 | DAKOCytomation | M7240 |
| **Secondary AB** | | | |
| donkey anti-goat IgG | Alexa Fluor 568 | Thermo Fisher Scientific | A-11057 |
| donkey anti-rabbit IgG | Alexa Fluor 488 | Jackson ImmunoResearch Lab | 711-546-152 |
| donkey anti-mouse IgG | Alexa Fluor 647 | Jackson ImmunoResearch Lab | 715-606-150 |
